# A cytosolic function of DNMT1 controls neuronal morphogenesis via microtubule regulation

**DOI:** 10.1101/2025.10.19.683279

**Authors:** Georg Pitschelatow, Junko Kurisu, Takumi Kawaue, Ke Zuo, Song Xie, Christoph Weber-Hamacher, Cathrin Bayer-Kaufmann, Jian Du, Philip Wolff, Shiori Nagayama, Claudia Palacios-Sanchez, Aylin Rogowski, Jana Egner-Walter, Paolo Ruggerone, Marc Spehr, Paolo Carloni, Mineko Kengaku, Geraldine Zimmer-Bensch

## Abstract

Proteins traditionally confined to a single cellular compartment are increasingly recognized to exert non-canonical functions in alternative domains. The DNA methyltransferase 1 (DNMT1), classically defined as the maintenance methyltransferase that preserves DNA methylation patterns during replication, exemplifies this versatility. Beyond its canonical role, DNMT1 is highly expressed in the developing and adult brain, where it contributes to transcriptional regulation in postmitotic neurons. Notably, cytoplasmic DNMT1 localization has been observed in neural cells, and emerging evidence links DNMT1 to mitochondrial function with implications for neurodegenerative disease, whereby the underlying functional mechanisms remain to be fully elucidated.

Here, we identify a previously unrecognized cytosolic function of DNMT1 in developing cortical excitatory neurons. Through genetic perturbation, proteomics, and high-resolution imaging, we show that DNMT1 regulates dendritic and axonal branching independently of its catalytic activity and nuclear localization. Instead, DNMT1 operates as a cytosolic scaffold interacting with the polarity regulator DOCK7 to modulate Rac1–STMN1 signaling, microtubule dynamics, and organelle trafficking. These findings expand the conceptual framework of DNMT1 from a genome guardian to a dual-compartment regulator that coordinates cytoskeletal remodeling and mitochondrial positioning. Beyond advancing our understanding of neuronal morphogenesis, this work provides mechanistic insight into how DNMT1 mutations may lead to neurodegenerative diseases.

## Introduction

It is increasingly appreciated that proteins traditionally confined to specific subcellular compartments can exert non-canonical functions in alternative cellular domains. This is particularly true for nuclear proteins with regulatory potential, whose spatial distribution and protein–protein interactions critically influence their functional repertoire^1^. One such protein is DNMT1, canonically recognized as the principal maintenance methyltransferase preserving cytosine methylation patterns during DNA replication^2^. Yet, recent studies suggest that DNMT1 also binds unmethylated DNA and regulates transcription in non-dividing cells, including postmitotic and fully differentiated neurons^3–5^.

DNMT1 is highly expressed in both the developing and adult brain^6,7^ and has been implicated in diverse neuronal functions extending beyond classical epigenetic maintenance^3,8,9,4^. In adult inhibitory interneurons of the cerebral cortex, DNMT1 regulates synaptic transmission by controlling endocytosis-related gene expression in a DNA methylation-dependent manner^3^. Moreover, in migrating preoptic area-derived cortical interneurons, DNMT1 influences cytoskeletal remodeling by interacting with histone modifications, particularly affecting the repressive histone mark H3K27me3^9,10^. Collectively, these findings highlight a broader functional spectrum of DNMT1 in the nervous system that is not limited to DNA methylation. Excitatory neurons, alongside inhibitory interneurons, form the principal neuronal populations of the cerebral cortex and arise in its proliferative zones. Recent work showed that DNMT1 contributes to the morphological maturation of cortical excitatory neurons during corticogenesis: *Emx1-Cre*–mediated *Dnmt1* deletion markedly increased dendritic branching complexity^4^. However, the molecular mechanisms underlying DNMT1-dependent cytoskeletal regulation in excitatory neurons remain unresolved, and indirect effects from astrocyte dysfunction cannot be excluded due to the broad expression pattern targeted by *Emx1-Cre*. Structurally, DNMT1 possesses the largest N-terminal regulatory domain among DNMT family members, containing multiple interaction motifs that mediate dynamic protein–protein interactions^11,12,13^. It binds to its well-established cofactor UHRF1, which couples DNMT1 activity to histone code interpretation at the replication fork, enabling maintenance of DNA methylation marks on newly synthesized daughter strands^14,15^. Moreover, DNMT1 can bind histone modifiers such as lysine-specific demethylase 1 (LSD1) and enhancer of zeste homolog 2 (EZH2), to modulate transcription independently of DNA methylation^10,16^. These properties position DNMT1 as a modular scaffold capable of engaging in distinct cellular processes. Supporting this view, disease-associated DNMT1 mutations cluster within interaction domains and correlate with distinct neurological phenotypes^17,18^, reinforcing the importance of its context-specific interactions.

While DNMT1 is classically described as a nuclear protein, early histological studies reported prominent cytoplasmic DNMT1 localization in adult neurons^19^, and subsequent work observed a similar relocalization in astrocytes following brain injury^20^. Moreover, DNMT1 has been recently shown to impact mitochondrial function^21,22^. Despite these observations, the functional significance of DNMT1’s extranuclear presence remains not fully understood. While it has been proposed that cytoplasmic retention merely regulates the nuclear pool^19^, potential cytosolic functions cannot be excluded.

In this study, we demonstrate a previously unrecognized cytosolic role of DNMT1 in shaping neuronal morphology during cortical development. Loss of DNMT1 increases dendritic and axonal branching independently of its catalytic activity and nuclear localization, implicating a cytoplasmic mechanism. We show that cytosolic DNMT1 associates with trafficking and cytoskeletal-related proteins, modulating microtubule stabilization via a DOCK7–Rac1– STMN1 pathway and mitochondrial trafficking. These findings challenge the view of DNMT1 as an exclusively nuclear methyltransferase, uncovering a dual-compartment role that links cytoskeletal remodeling and organelle trafficking in developing neurons.

## Results

### *Dnmt1* knockdown increases dendritic complexity in developing excitatory cortical neurons in vivo

DNMT1 is expressed in both inhibitory and excitatory neurons of the developing cortex^4,5,9^. Conditional *Dnmt1* deletion in excitatory neurons (*Emx1-Cre*) was previously shown to increase dendritic branching complexity^4^, suggesting a regulatory role of DNMT1 in neuronal morphogenesis. However, the molecular mechanisms underlying the DNMT1-dependent cytoskeletal regulation in excitatory neurons remain unknown. Moreover, because *Emx1-Cre*– mediated deletion also targets later-born astrocytes, indirect, astrocyte-mediated effects cannot be excluded in these earlier in vivo studies.

To directly assess whether DNMT1 regulates embryonic cortical neuron morphology, we employed a CRISPR interference (CRISPRi) system to repress *Dnmt1* expression via in utero electroporation of a pX458-CAG-dCas9KRAB-2A-Dnmt1*-*GFP plasmid (CRISPRi-Dnmt1). To validate CRISPRi-Dnmt1 construct efficiency, we transfected N2a cells and performed immunohistochemical staining against DNMT1. Signal intensities, quantified as integrated densities (ID) and normalized to the DAPI signal, revealed a significant reduction in DNMT1 protein level in CRISPRi-Dnmt1-transfected cells, compared to control transfections (pX458-CAG-dCas9KRAB-2A-GFP plasmid; Supplementary Fig. 1a-c).

For in vivo knockdown, either CRISPRi-Dnmt1 or control plasmids were targeted to the cortical proliferative zones of E12.5 embryos via electroporation, followed by analysis at P5. Robust GFP expression was detected in the cerebral cortex in 300 µm vibratome sections, co-stained with DAPI and for CTIP2 to label deep-layer cortical neurons^23^ (Fig. 1a, b). 3D reconstruction of CTIP2⁺ neurons using Imaris (Fig. 1c, d) revealed a significant increase in the number of basal dendrites and arbor complexity in *Dnmt1*-depleted neurons relative to controls (Fig. 1e), with no obvious defects in laminar positioning. These data suggest that DNMT1 cell-autonomously restrains dendritic outgrowth, in line with earlier observations by Hutnick et al. (2009)^4^.

**Figure 1:**
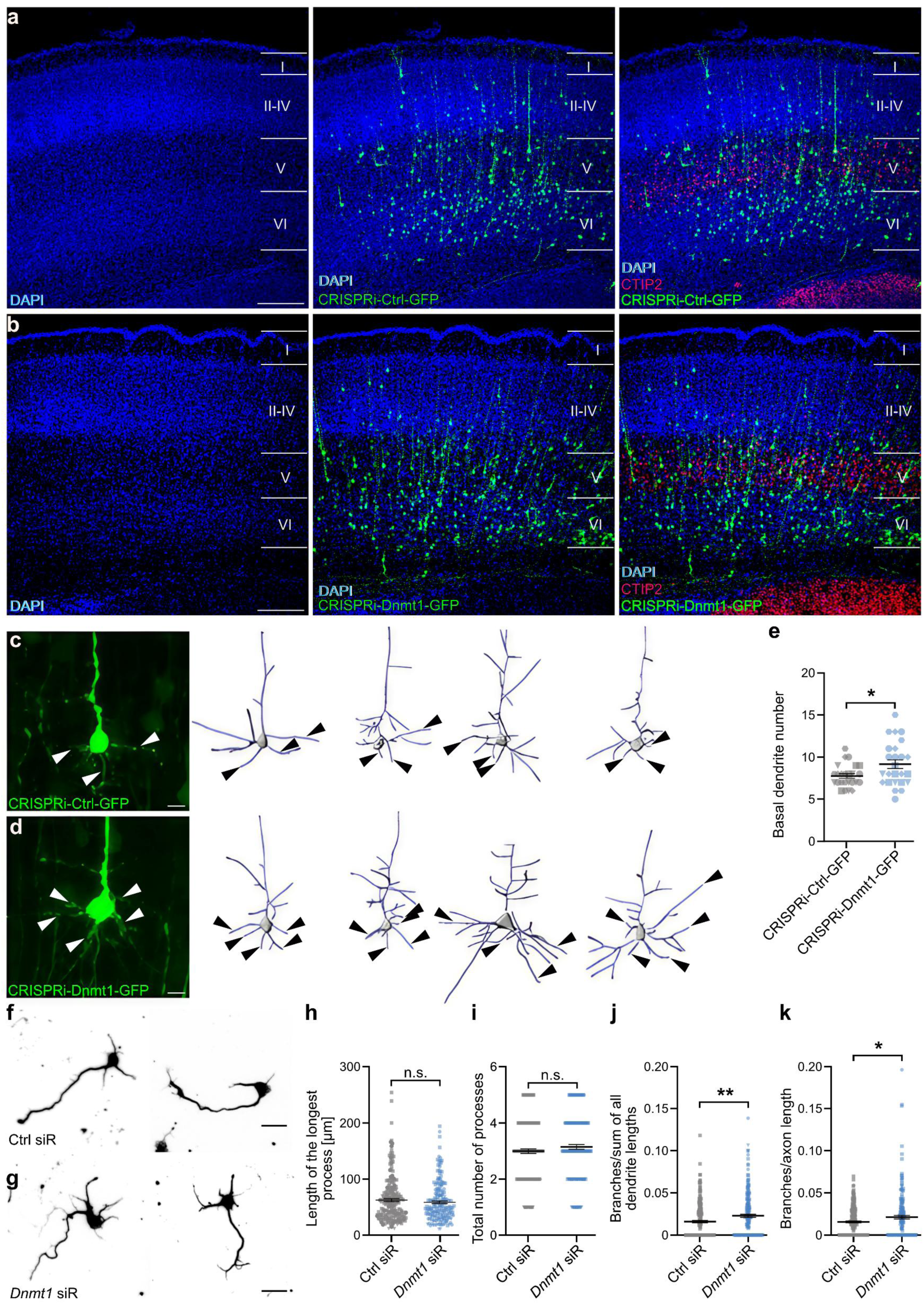
Repression of *Dnmt1* expression by the CRISPRi-Dnmt1-GFP construct or *Dnmt1* siRNA increases the basal dendrite number and branching in cortical neurons in vivo and in vitro. (**a, b**) Exemplary brain sections (300 µm) from P5 brains of pups, which were intrauterine electroporated at E12.5 with either CRISPRi-Ctrl-GFP (**a**) or CRISPRi-Dnmt1-GFP (**b**), followed by immunostaining for GFP (CRISPRi-Ctrl-GFP/CRISPRi-Dnmt1-GFP, green), CTIP2 (red), and DAPI (blue). Distinct cortical layers are labeled from I to VI. Scale bars: 200 µm. (**c, d**) Exemplary images (left) and 3D-reconstructions (right) of exemplary GFP-positive laver V cortical neurons from the CRISPRi-Ctrl-GFP (**c**) or the CRISPRi-Dnmt1-GFP (**d**) conditions at P5. The white/black arrow heads highlight basal dendrites. Scale bars: 10 µm. (**e**) Analysis of the absolute basal dendrite number. n (CRISPRi-Ctrl-GFP) = 25 cells; n (CRISPRi-Dnmt1-GFP) = 25 cells. N = 4 experiments. Welch’s t-test, * p < 0.05. (**f, g**) Inverted microphotographs of exemplary βIII-tubulin immunocytochemically stained cortical neurons (E14.5 + 2 DIV) previously transfected with control (**f**) or *Dnmt1* (**g**) siRNA at 1 DIV for 24 h. Scale bars: 20 µm. (**h-k**) Quantitative analysis of the length of the longest process (**h**), the number of processes (**i**), the branches normalized to the sum of the lengths of all processes (except the axon) presumably representing dendrites (**j**), and the branches per longest process length likely representing the axon (**k**). n (Ctrl siR) = 207 cells; n (*Dnmt1* siR) = 197 cells. N = 3 experiments. Two-tailed Student’s t-test, * p < 0.05, ** p < 0.01. Data are presented as mean ± SEM.

These in vivo findings were reproduced in vitro in E14.5 primary cortical neurons transfected with the CRISPRi-Dnmt1 construct (Supplementary Fig. 1d-h). A comparable phenotype was also evident in CRISPRi-Dnmt1-transfected N2a cells (Supplementary Fig. 1i-m). However, given the low transfection efficiency of plasmid DNA in primary neurons (∼10–20%, data not shown), we additionally employed siRNA-mediated knockdown of *Dnmt1* to increase the number of targeted cells (transfection efficiency > 50 %, data not shown). A detailed morphological analysis revealed that *Dnmt1* siRNA treatment of cortical neurons (E14.5 + 1 DIV) led to a significant increase in both axonal and dendritic branching compared to control oligonucleotides after 24 hours (Fig. 1f-k), with axons identified via SMI immunostaining (Supplementary Fig. 1n, o; siRNA knockdown efficiency was already confirmed previously^9^). A similar increase in neurite complexity was observed in E14.5 cortical neurons treated with *Dnmt1* siRNA at 3 DIV and analyzed at 5 DIV, a time point chosen to capture more advanced morphological features (Supplementary Fig. 2a-e). Furthermore, P0 cortical neurons transfected with *Dnmt1* or control siRNA at 3 DIV exhibited increased axonal branching upon analysis at 5 DIV (Supplementary Fig. 2f-j). These findings indicate that the branching-restricting effect of DNMT1 is conserved across developmental stages and neuronal models.

### Branching regulation is independent of DNMT1’s catalytic activity

As described, CRISPRi and siRNA-based knockdown of *Dnmt1* in cortical neurons and N2a cells increased neurite branching (Fig. 1, Supplementary Fig. 1, 2). Next, we aimed to assess the molecular mechanisms of DNMT1-dependent cytoskeletal regulation. We used the small molecule inhibitor RG108 (*N-Phthalyl-L-tryptophan)*, known to selectively block the catalytic activity of DNA methyltransferases by binding to its active site without incorporating into DNA, thereby reducing global DNA methylation levels^24^. We first confirmed that RG108 treatment significantly reduced global 5mC levels by immunostaining in N2a cells (Supplementary Fig. 3a-c). However, RG108 treatment did not mimic the morphological phenotype of *Dnmt1* knockdown in cortical neurons (E14.5 + 2 DIV; Supplementary Fig. 3d-h), suggesting that DNMT1’s role in controlling neuronal morphology is independent of its catalytic activity.

To more directly assess the non-canonical role of DNMT1, we generated a series of DNMT1 mutant constructs resistant to *Dnmt1* siRNA through silent point mutations. These included CAG-mNG-Dnmt1-WT (wildtype), ΔCat (lacking the catalytic domain), ΔpCat (bearing a catalytic point mutation), and ΔNLS (lacking the nuclear localization sequence), all tagged with mNeonGreen (mNG) and a V5 epitope (Fig. 2a). Following sequence validation (Supplementary Table 2), constructs were transfected into N2a cells, showing the expected subcellular localization. While WT, ΔCat, and ΔpCat were localized predominantly in the nucleus, ΔNLS was restricted to the cytoplasm (Fig. 2b). Comparable findings were observed 24 hours post-transfection in cortical neurons (E14.5 + 2 DIV, Fig. 2c).

**Figure 2:**
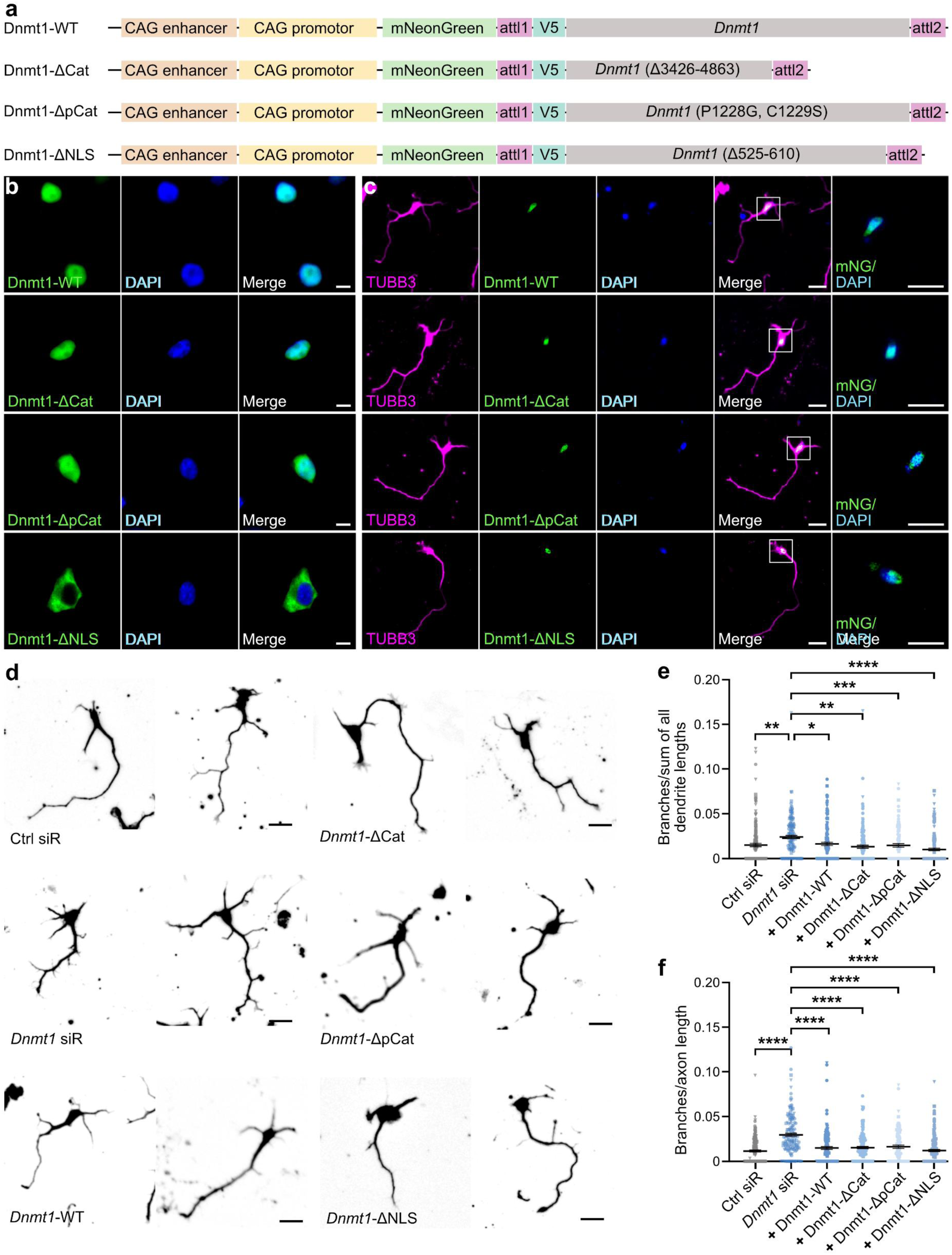
Mutant Dnmt1 constructs lacking catalytic activity or nuclear localization sequence restore branching in *Dnmt1*-depleted neurons and DNMT1 is localized in the cytosol. (**a**) Scheme of the different generated Dnmt1 expression constructs demonstrating all their relevant functional domains, tags, and promotor. (**b**) Representative fluorescence microphotographs of N2a cells expressing siRNA resistant mNG-DNMT1 (WT, ΔCat, ΔpCat, ΔNLS) plasmids (green) 24 h post co-transfection with *Dnmt1* siRNA and followed by immunostaining with DAPI (blue). Scale bars: 10 µm. (**c**) Representative microphotograph of cortical neurons (E14.5 + 2 DIV) co-transfected with *Dnmt1* siRNA and the different Dnmt1 plasmids (green) at 1 DIV for 24 h, which were later immunocytochemically stained for βIII-tubulin (TUBB3, magenta) and DAPI (blue). The white squares represent the respective magnification of the merge. Scale bars: 20 µm. (**d**) Exemplary inverted microphotographs of βIII-tubulin-stained cortical neurons (E14.5 + 2 DIV) under the same co-transfection conditions. Scale bars: 20 µm. (**e, f**) Analysis of the branches normalized to the longest process length presumably representing axons (**f**) and branches per length cumulated across all remaining process lengths likely representing dendrites (**e**). “+” indicates neurons, which were also successfully co-transfected with *Dnmt1* siRNA for the downregulation of the endogenous *Dnmt1* expression in addition to the Dnmt1 plasmids. n (D1-WT) = 141 cells; (D1-ΔCat) = 136 cells; n (D1-ΔpCat) = 98 cells; n (D1-ΔNLS) = 150 cells; n (Ctrl siR) = 153 cells; n (*Dnmt1* siR) = 146 cells. N = 3 experiments. One-way ANOVA with Tukey’s post-hoc multiple comparison test, * p < 0.05, ** p < 0.01, *** p < 0.001, **** p < 0.0001. Data are presented as mean ± SEM.

For functional analysis, E14.5 + 1 DIV cortical neurons were co-transfected with *Dnmt1* siRNA to knockdown endogenous *Dnmt1*, along with the respective Dnmt1 rescue plasmids. Reference conditions included neurons transfected with either control siRNA or *Dnmt1* siRNA alone. Morphological parameters were quantified in neurons co-labeled for mNG and siRNA, or siRNA alone as reference. *Dnmt1* siRNA treatment increased dendritic and axonal branching (Fig. 2d-f), consistent with previous experiments (Fig. 1f-k). Importantly, co-expression of any Dnmt1 mutant construct—including catalytically inactive and cytosolic variants—restored axonal and dendritic branching to control levels (Fig. 2d-f). These data indicate that DNMT1 modulates the morphology of developing cortical neurons independently of its DNA methyltransferase activity or even independent of its nuclear localization. This suggests a non-canonical, and so far unreported, cytosolic function of DNMT1 in regulating axonal and dendritic branching in developing cortical neurons.

### DNMT1 is partially localized in the cytosol and interacts with cytoskeletal and trafficking proteins

Although DNMT1 is classically described as a nuclear maintenance methyltransferase, recent evidence suggests that DNMT1 may also localize to the cytoplasm in neurons and glial cells^19,20^. These studies are in line with our own rescue experiments using Dnmt1 mutants, where the ΔNLS construct—lacking the nuclear localization sequence—was able to rescue the morphological phenotype induced by *Dnmt1* knockdown, despite being confined to the cytosol (Fig. 2). These findings raised the hypothesis that DNMT1 might exert functional, non-canonical roles in the cytoplasm. To investigate the subcellular localization of DNMT1 in more detail, we performed immunolabeled DNMT1 in E14.5 + 2 DIV cortical neurons and N2a cells (after 48 h), visualizing its distribution using high-resolution STED microscopy. DNMT1 was clearly detected within the nucleus, overlapping with DAPI staining (Fig. 3a, b). However, distinct DNMT1-positive puncta also became evident in the cytosolic compartment in both cell types (Fig. 3a, b), suggesting extranuclear localization. Additional evidence for cytosolic DNMT1 was provided by subcellular fractionation and Western blot analysis of N2a cells, which confirmed DNMT1 presence in cytosolic fractions (Supplementary Fig. 3i).

**Figure 3:**
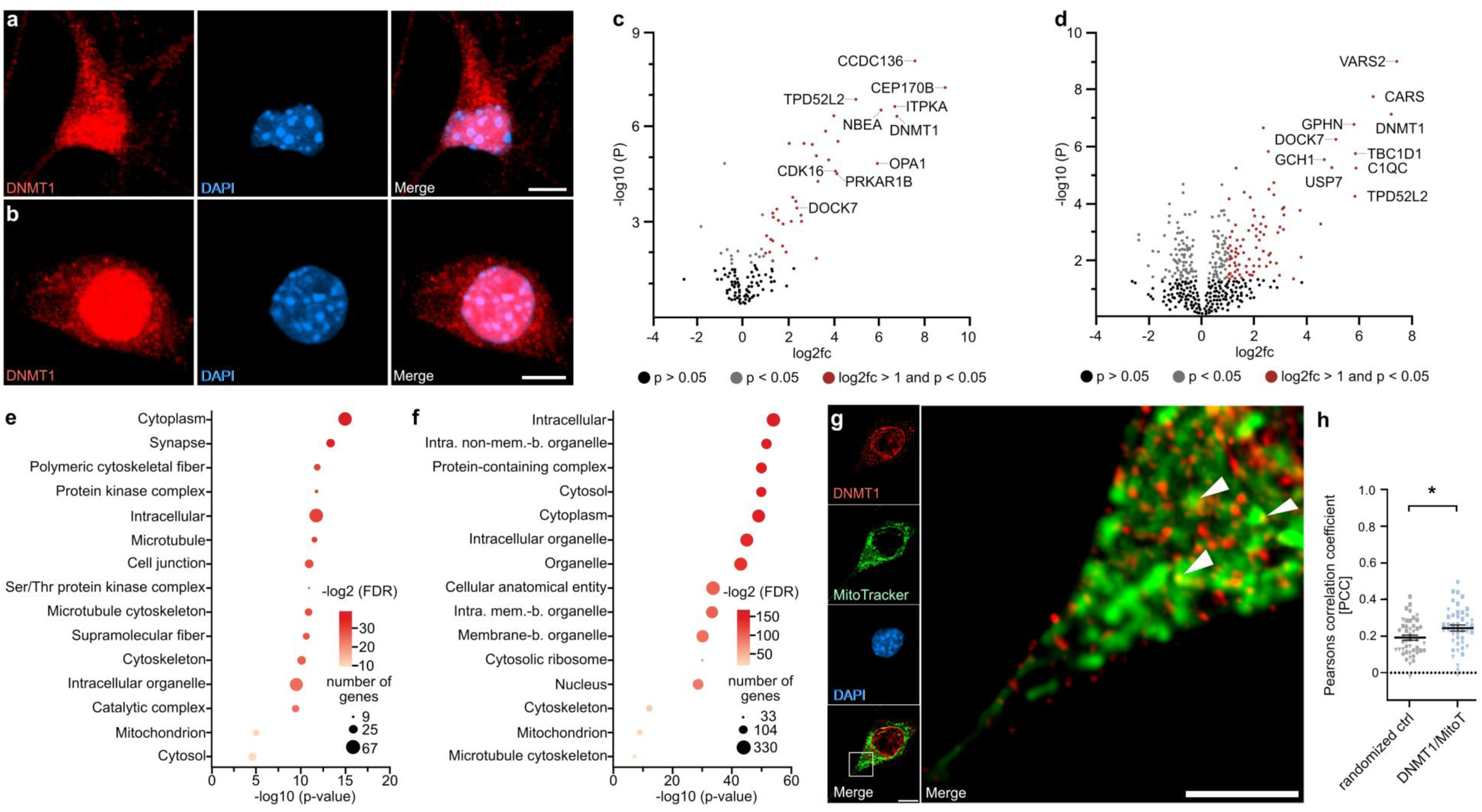
DNMT1 is partially localized in the cytosol and interacts with cytosolic proteins. **(a, b)** Immunocytochemical staining for DNMT1 (red) and DAPI (blue) in a cortical neuron (E14.5 + 2 DIV) **(a)** and an N2a cell **(b)**, visualized by high-resolution STED microscopy. Scale bars: 5 µm. **(c, d)** Volcano plots showing proteins co-immunoprecipitated with DNMT1 from murine brain tissue lysates (3.5 months; c, Supplementary Data 1) and N2a cell lysates (d, Supplementary Data 2), identified by mass spectrometry. N = 4 biological replicates for each condition. **(e, f)** Gene Ontology (GO) enrichment analysis of DNMT1 co-immunoprecipitated proteins from murine brain tissue **(e)** and N2a cells **(f)**, performed using the STRING database and ShinyGO (v0.82). **(g)** Representative STED micrograph of a fixed N2a cell cultured for 48 h, labeled with MitoTracker™ Deep Red FM (shown in false color green), DAPI (blue), and an antibody against DNMT1 (red). Scale bar: 5 µm. The white box indicates the magnified region shown in the inset (Scale bar: 2 µm). Nuclear signals were computationally removed to enhance visualization of the cytosolic compartment for further analysis. **(h)** Quantification of cytosolic colocalization between DNMT1 and mitochondria by Pearson’s correlation coefficient (PCC) compared to a rotated randomized control. n (ROIs) = 45; N (cells) = 15. Two-tailed Student’s t-test, p < 0.05, p < 0.01, p < 0.001, p < 0.0001.

Given DNMT1’s large N-terminal domain, known for mediating protein–protein interactions^11,12,13^, we hypothesized that DNMT1 may engage in cytoplasmic processes underlying neurite branching regulation, such as cytoskeletal remodeling. To gain mechanistic insight into which cytosolic processes DNMT1 is involved in, we next set out to verify and characterize its interaction profile. To this end, we conducted co-immunoprecipitation on lysates from cortical tissue and N2a cells, followed by LC-MS/MS analysis, which revealed DNMT1 interactors not only in the nucleus but also in the cytoplasm (Fig. 3c-f, Supplementary Data 1, 2). Of note, many detected DNMT1 interacting proteins were also identified in a recent study in human cells^22^. GO analysis of our dataset highlighted partners involved in microtubule organization and trafficking, including DOCK7, MAP2, TPD52L2, KIF5B, and KIFC1^25–28, 29^ (Fig. 3c, d, Supplementary Fig. 4a, b). These interactions place DNMT1 in a cytosolic hub at the microtubule–organelle interface, where it could couple microtubule stabilization factors (DOCK7, MAP2) with transport machinery (KIF5B/KIFC1) and vesicle regulators (TPD52L2). MAP2, one prominent interactor, binds microtubules via its tubulin-binding domain and regulates neurite outgrowth, dynamics, and organelle transport^30^. MAP2 can also bridge mitochondria to microtubules^31^. Interestingly, and in line with other studies^21,22^, mitochondrial proteins were also enriched in the DNMT1 interactome (Fig. 3e, f). Moreover, partial DNMT1colocalization with mitochondria was confirmed in N2a cells (Fig. 3g, h; Pearson’s correlation coefficient (PCC) ∼0.21; Supplementary Fig. 4b, c). In contrast, no overlap with ER or lysosomes was detected (Supplementary Fig. 4d, e). Thus, DNMT1 is positioned to influence cytoskeleton-mitochondria interactions and trafficking.

### DNMT1 modulates mitochondrial trafficking without affecting mitochondrial function

Mitochondria are trafficked along microtubules to branch sites, providing local ATP essential for axonal and dendritic growth^32,33,34^. Given DNMT1’s cytosolic role in branching regulation, its partial co-localization with mitochondria, and its interaction with mitochondrial proteins as well as cytoskeletal-related proteins that interact with mitochondria (Fig. 3, Supplementary Fig. 4), we hypothesized that it may affect mitochondria trafficking.

To investigate the role of DNMT1 in mitochondrial trafficking, P0 primary cortical neurons were prepared and co-transfected at DIV 3 with a mitochondria-targeted GFP construct (MT-GFP) and either *Dnmt1* siRNA (together with Alexa Fluor™ 555-conjugated control siRNA) or control siRNA alone (Fig. 4a). As reported before, this experimental design also revealed increased axonal branching upon *Dnmt1* depletion (Supplementary Figure 2f-j) and allowed monitoring organelle dynamics due to the extended length of axons.

**Figure 4:**
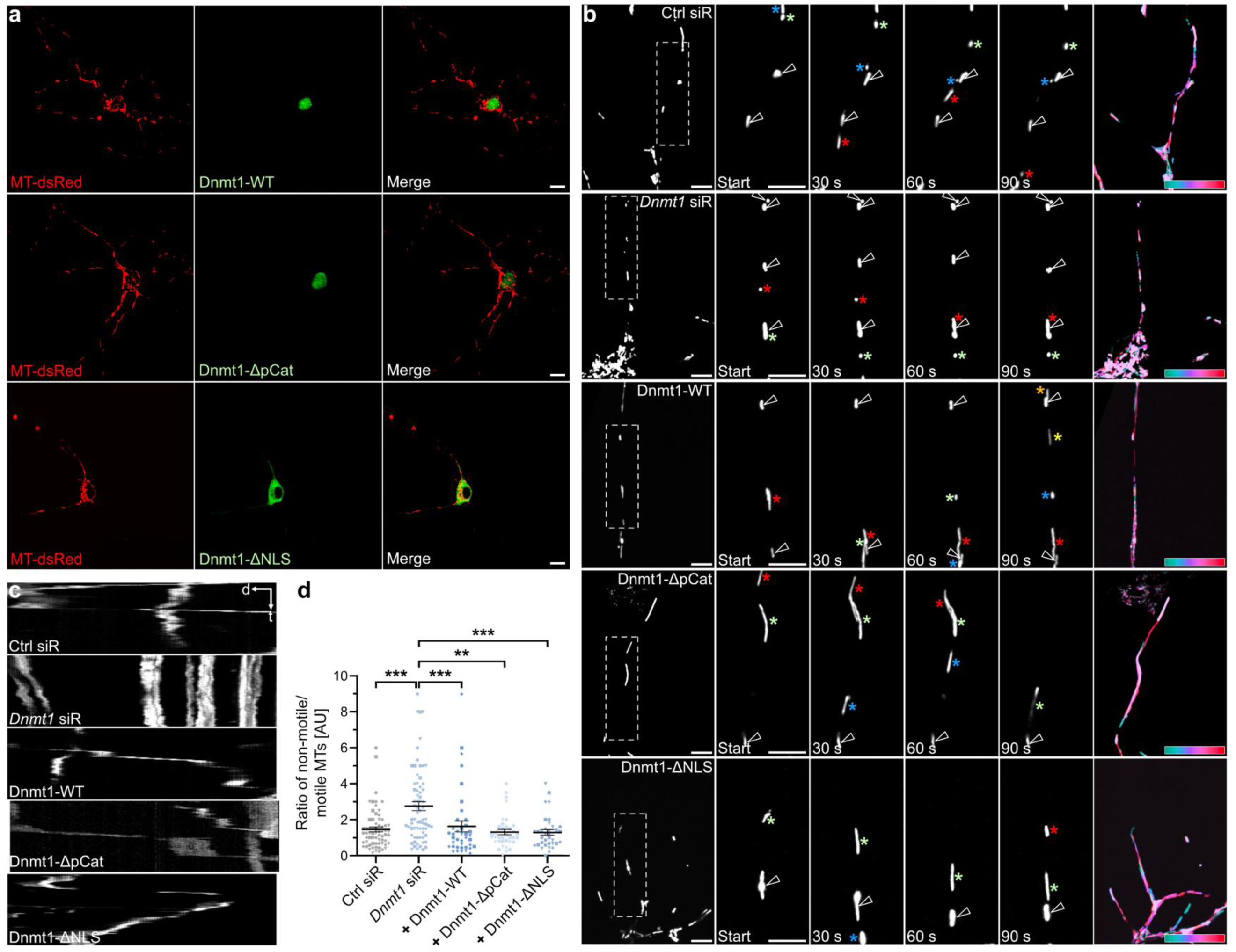
Expression of Dnmt1 plasmid constructs rescue the *Dnmt1* siRNA-induced increase in non-motile mitochondria. **(a)** Representative images of cortical neurons (P0 + 4 DIV) co-transfected at DIV 3 with MT-dsRed (red) and the indicated Dnmt1 plasmid constructs (green) for 24 h. Scale bars: 10 µm. **(b)** Exemplary tracks of MT-dsRed–labeled mitochondria in cortical neurons under the respective Dnmt1 expression conditions. Non-motile mitochondria are indicated by white arrowheads; motile mitochondria are marked with colored stars (each color corresponds to one mitochondrion). White boxes in the overview images (first panel) indicate magnified regions shown across time frames. The final frame (sixth panel) displays temporal color coding of mitochondrial trajectories. Scale bars: 5 µm. **(c)** Representative kymographs illustrating mitochondrial motility under the respective conditions. **(d)** Quantification of the ratio of non-motile to motile mitochondria in cortical neurons (P0 + 4 DIV). “+” indicates neurons co-transfected with *Dnmt1* siRNA to downregulate endogenous *Dnmt1* expression in addition to the respective Dnmt1 expression constructs. N (D1-WT) = 39 ROIs; n (D1-ΔpCat) = 36 ROIs; n (D1-ΔNLS) = 35 ROIs; n (Ctrl siR) = 71 ROIs; n (*Dnmt1* siR) = 75 ROIs. N = 4-7 independent experiments. Kruskal–Wallis test followed by Dunn’s post hoc multiple comparison test, p < 0.01, p < 0.001. Data represent mean ± SEM.

Live-cell imaging at DIV 4 revealed that *Dnmt1* knockdown significantly increased the fraction of non-motile mitochondria (Fig. 4b-d, Supplementary Movies M1 and M2), indicating altered trafficking and/or docking dynamics. Of note, mitochondrial size remained unchanged (Supplementary Fig. 5a-c), ruling out a causal role of altered organelle fission/fusion.

To test whether DNMT1’s non-canonical, cytosolic functions regulate mitochondria trafficking, we performed rescue experiments using DNMT1 mutant constructs (WT, ΔpCat, ΔNLS) alongside *Dnmt1* siRNA and mitochondria-targeted dsRed (MT-dsRed). Co-expression of DNMT1-WT rescued the reduction in mitochondrial motility triggered by *Dnmt1* siRNA (Fig. 4c, d, Supplementary Movies M3). Importantly, co-expression of ΔpCat and ΔNLS with *Dnmt1* siRNA also restored the ratio of non-motile/motile mitochondria (Fig. 4c, d, Supplementary Movies M4 and M5), supporting a DNA methylation-independent, cytoplasmic mechanism underlying DNMT1-dependent regulation of mitochondrial transport.

We further assessed whether DNMT1 influences mitochondrial activity using tetramethylrhodamine ethyl ester (TMRE) labeling in living N2a cells. TMRE accumulates in active mitochondria in proportion to their membrane potential, serving as an indirect readout of mitochondrial function^35^. Carbonyl cyanide-4-(trifluoromethoxy)phenylhydrazone (FCCP)-treated cells served as positive controls, showing the expected TMRE signal reduction due to membrane depolarization^36^ (Supplementary Fig. 5d-f). FCCP collapses the proton gradient across the inner mitochondrial membrane, thereby dissipating the mitochondrial membrane potential^37^. In contrast to FCCP treatment, *Dnmt1* siRNA application had no significant effect on TMRE signal intensity compared to control oligos, suggesting that DNMT1 does not affect the mitochondrial membrane potential (Supplementary Fig. 5g, h).

In sum, our results show that DNMT1 depletion disrupts mitochondrial trafficking, while mitochondrial activity and morphology remain unaffected. Since efficient mitochondrial transport depends on stable microtubules, and DNMT1 interacts with microtubule-regulating proteins, we next asked whether DNMT1 affects microtubule stability.

### DNMT1 and DOCK7 physically interact and share a role in neuronal morphology regulation

DOCK7 regulates microtubules by activating Rac GTPases, which then promote phosphorylation and inactivation of the microtubule-destabilizing protein stathmin/Op18, thereby stabilizing microtubules^26^. DOCK7 was identified as a DNMT1 interactor in both N2a cells and cortical tissue (Fig. 3c, d). Given DOCK7’s known function in regulating neuronal polarity via the Rac1-STMN1 axis^26^, we examined its interaction with DNMT1 in more detail. Co-immunoprecipitation in cortical lysates confirmed a physical interaction between DNMT1 and DOCK7 (Fig. 5a, b). Moreover, high-resolution STED microscopy of double immunostaining against DOCK7 and DNMT1 in N2a cells revealed partial colocalization of both proteins in the cytoplasm (Fig. 5c-e).

**Figure 5:**
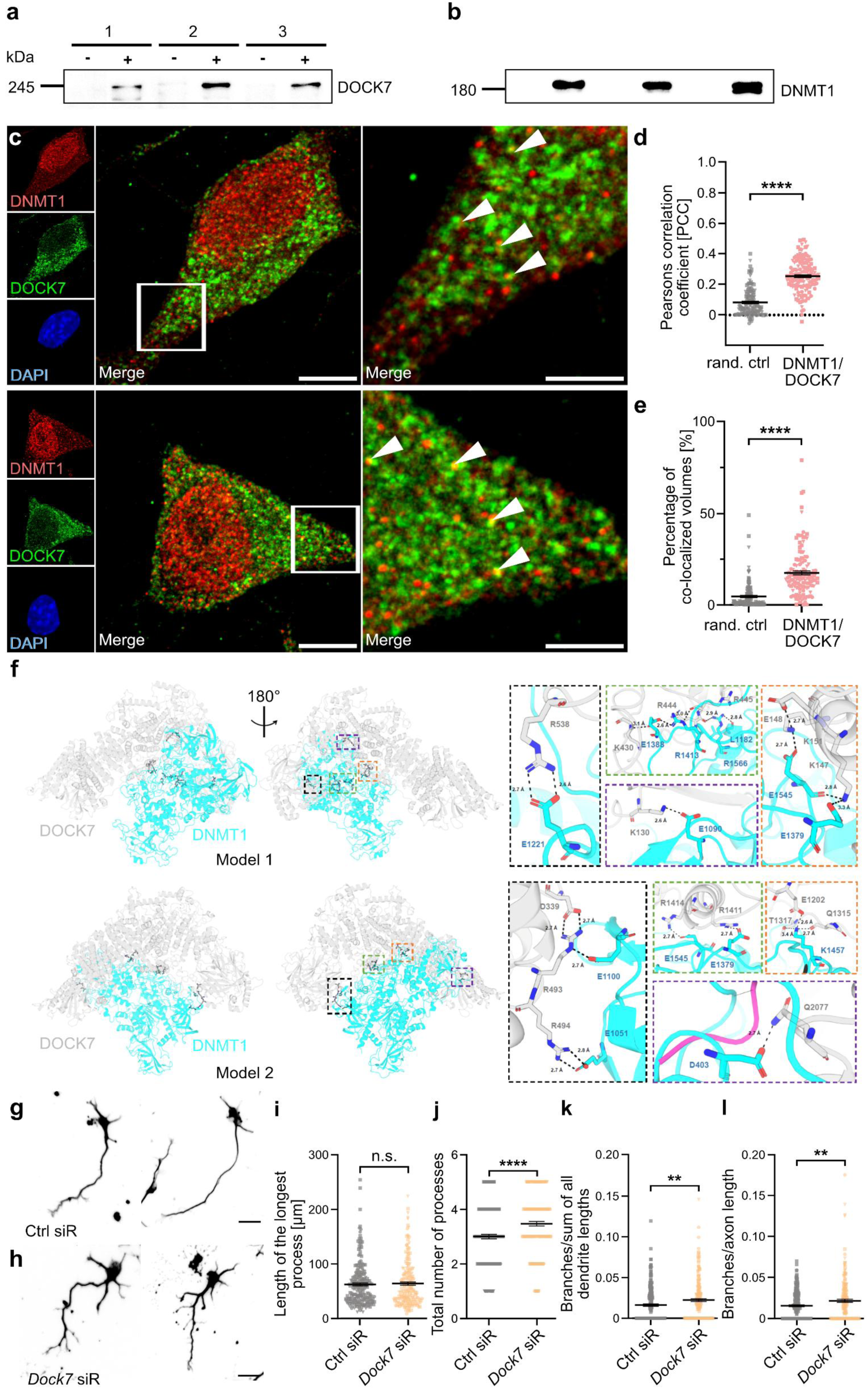
DOCK7 interacts with DNMT1 and knockdown of *Dock7* increases neuronal branching. (**a, b**) DOCK7 (TMW = 239 kDa, **a**) and DNMT1 (TMW = 183 kDa, **b**) protein interaction validated by DNMT1 co-immunoprecipitation and Western blot analysis by using a specific antibody against DOCK7 and DNMT1 in E14.5 cortical lysates. N = 3 experiments. + = DNMT1-antibody pulldown, - = IgG-antibody pulldown. (**c**) Representative images of N2a cells (grown for 48 h) co-stained with conjugated antibodies directed against DNMT1 (red) and DOCK7 (green), additionally stained with DAPI (blue), and captured by high-resolution STED microscopy. Scale bars: 5 µm. The white squares depict the magnification of the merge. Scale bars: 2 µm. Overlapping volumes are highlighted by white arrowheads. (**d, e**) Analysis of cytosolic colocalization between DNMT1 and DOCK7 by Pearson’s correlation coefficient (PCC) (**d**) and the percentage of colocalized volumes (**e**) in comparison to the corresponding rotated and randomized controls. n (ROIs) = 133; N (cells) = 45. (**f**) Predicted interaction between DNMT1 and DOCK7 using MD simulations and docking methods (left figure). Specific hot spot residues for the interaction between DNMT1 and DOCK7 are represented in the right figure. Hydrogen bonding and salt bridge interactions are shown as black dashes. (**g, h**) Inverted microphotographs of exemplary βIII-tubulin immunocytochemically stained cortical neurons (E14.5 + 2 DIV) transfected with control (**g**) or *Dock7* (**h**) siRNA at 1 DIV for 24 h. Scale bars: 20 µm. (**i-l**) Analysis of morphological parameters, such as the length of the longest process (**i**), the number of processes (**j**), the branches per length summed across all processes likely representing dendrites (**k**), and the branches normalized to the longest process length likely representing axons (**l**). n (Ctrl siR) = 207 cells; n (*Dock7* siR) = 198 cells. N = 3 experiments. Two-tailed Student’s t-test, * p < 0.05, ** p < 0.01, **** p < 0.0001. Data are presented as mean ± SEM.

In line with that, molecular Dynamics (MD) simulations and multiple docking approaches, including ClusPro (energy-based sampling and clustering global docking)^38^, LZerD (3D Zernike descriptors-driven free docking)^39^, RosettaDock (Monte Carlo docking with side-chain repacking)^40^, and GRAMM (shape complementarity-based global docking) ^41^, predicted two structural models of the DNMT1/DOCK7 complex (Models 1 and 2 in Fig. 5f, left). Both feature a large complementary interface between DNMT1 and DOCK7. The hot spot interfacial residues involve DNMT1 residues E1379, R1413, K1457, and R1566, and DOCK7 residues K130, K147, K151, K430, R493, R494, R538, T1317, R1411, R1414, and Q2077 (Fig. 5f, right). The interface is stabilized by contacts between these amino acids, primarily within the catalytic domain of DNMT1 and the region of DOCK7 interacting with it (Fig. 5f, right).

Notably, siRNA-mediated depletion of *Dock7* phenocopied *Dnmt1* knockdown, leading to increased axonal and dendritic branching in primary cortical neurons (Fig. 5g-l). In N2a cells, neurite branching was likewise elevated upon *Dock7* depletion (Supplementary Fig. 6a-e). Together with the observed interaction between DNMT1 and DOCK7, these data indicate a concerted action of both proteins in regulating neuronal morphology.

### DNMT1 and DOCK7 both influence mitochondrial trafficking and positioning

Consistent with its cytosolic localization, DOCK7 strongly colocalized with mitochondria (PCC ∼0.7) (Fig. 6a, b). To investigate whether DOCK7 also influences mitochondrial trafficking, primary cortical neurons were co-transfected with a mitochondria-targeted GFP construct (MT-GFP) and either *Dock7* siRNA or control siRNA. As an internal reference, *Dnmt1* siRNA was included in parallel to replicate the previously observed trafficking phenotype within the same set of experiments (Fig. 6c, d; Supplementary Movies M6, M7 and M8).

**Figure 6:**
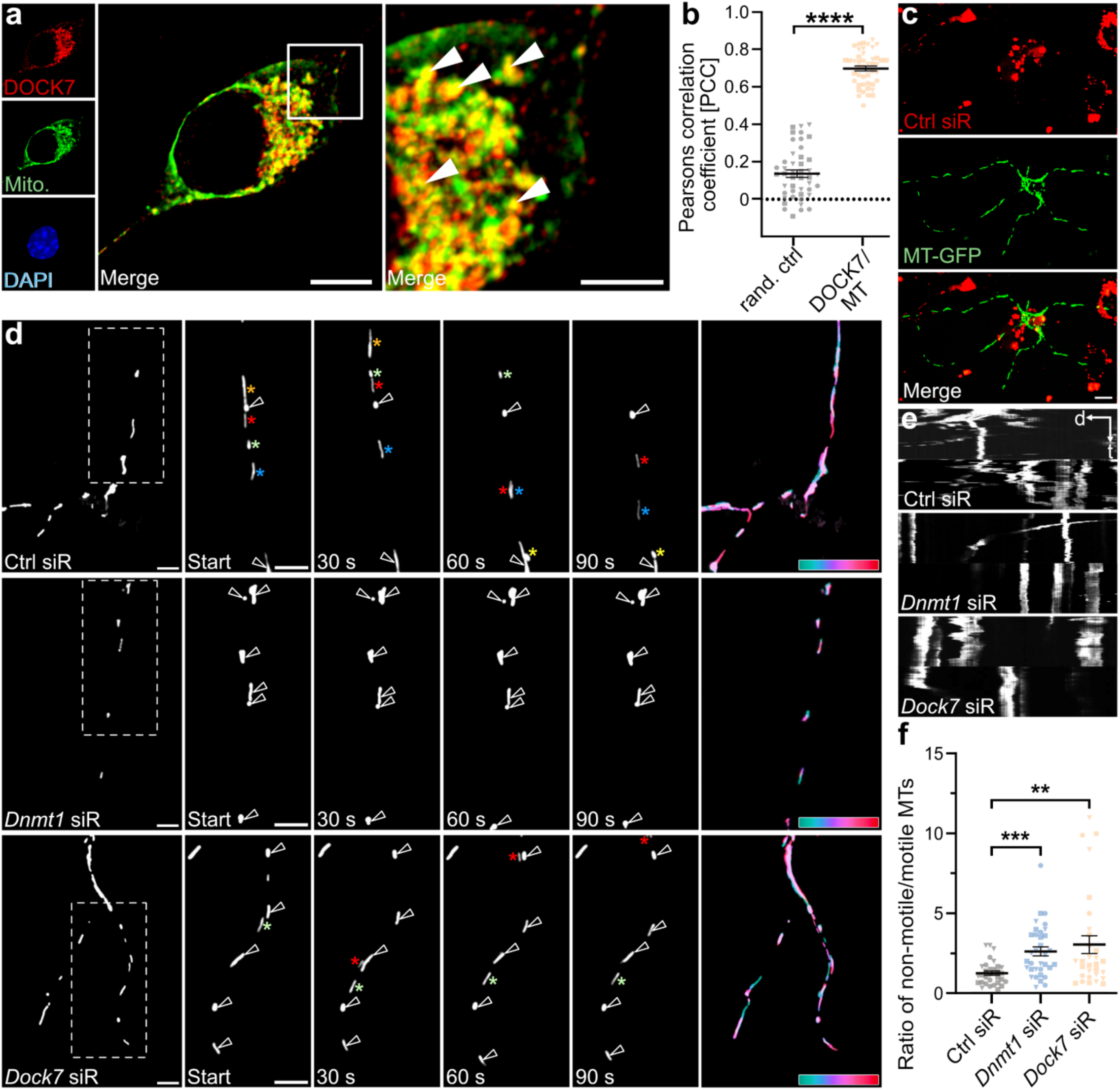
DOCK7 colocalizes with mitochondria, and its depletion increases the fraction of non-motile mitochondria. **(a)** Representative STED micrographs of a fixed N2a cell cultured for 48 h, labeled with MitoTracker™ Deep Red FM (Mito., false color green), DAPI (blue), and an antibody against DOCK7 (red). Scale bar: 5 µm. The white box indicates the magnified region shown in the merge (Scale bar: 2 µm). Nuclear signals were computationally removed to improve visualization of the cytosolic compartment. **(b)** Quantification of cytosolic colocalization between DOCK7 and mitochondria by Pearson’s correlation coefficient (PCC) compared to a rotated randomized control. n (ROIs) = 45; N (cells) = 15. Two-tailed Student’s *t*-test, p < 0.0001. **(c)** Representative images of cortical neurons (P0 + 4 DIV) co-transfected at DIV 3 with Alexa Fluor™ 555–labeled control siRNA (red) and MT-GFP plasmid (green) and imaged 24 h post-transfection. Scale bar: 10 µm. **(d)** Exemplary tracking of MT-GFP–labeled mitochondria in cortical neurons following control, *Dnmt1*, or *Dock7* siRNA treatment. Non-motile mitochondria are marked by white arrowheads, motile mitochondria by colored stars (each color indicates one mitochondrion). White boxes in overview images (first panel) mark the magnified regions shown over time. The last frame (sixth panel) depicts temporally color-coded mitochondrial trajectories. Scale bars: 5 µm. **(e)** Representative kymographs illustrating mitochondrial motility under the respective conditions. **(f)** Quantification of the ratio of non-motile to motile mitochondria in cortical neurons (P0 + 4 DIV) following control, *Dnmt1*, or *Dock7* siRNA-mediated knockdown for 24 h at DIV 3. N (Ctrl siR) = 35 ROIs; n (*Dnmt1* siR) = 39 ROIs; n (*Dock7* siR) = 33 ROIs. N = 3 independent experiments. Kruskal–Wallis test followed by Dunn’s post hoc multiple comparison test, p < 0.01, *p* < 0.001. Data are shown as mean ± SEM.

Live-cell imaging revealed that *Dock7* depletion, like *Dnmt1* knockdown, significantly increased the fraction of non-motile mitochondria (Fig. 6e, f, Supplementary Movies M6, M7 and M8), while mitochondrial size was not affected, proposing no effect on mitochondrial fission (Supplementary Figure 6f-h). Moreover, mitochondrial activity was likewise unaffected by *Dock7* knockdown (Supplementary Fig. 6i, j). In sum, DOCK7 and DNMT1 interact and elicit similar effects on neuronal morphology and mitochondria trafficking velocity.

Mitochondrial localization to prospective branch sites has been implicated in axonal and dendritic branching, with some studies associating mitochondrial accumulation with branch promotion, whereas others report inhibitory effects during early dendritic morphogenesis^42,43^. Building on these observations and considering our findings of impaired mitochondrial transport dynamics upon *Dnmt1* and *Dock7* knockdown, we next asked whether mitochondrial accumulation at branch initiation sites is impaired.

To this end, we performed live-cell STED imaging of MitoTracker-labeled N2a cells over 5 h (Fig. 7a; Supplementary Movies M9-M11). In control cells, mitochondrial density steadily increased at prospective branch points prior to outgrowth in control conditions, in line with what was reported in the literature^42^. In contrast, both *Dnmt1* and *Dock7* knockdown blunted this local accumulation, with both treatments resulting in reduced enrichment of the MitoTracker signal (Fig. 7a, b; Supplementary Movies M9-M11). That both manipulations promote neurite branch formation was also reproduced in this live cell imaging setting, reflected by increased frequency of branch initiation and shortened branch formation time (Fig. 7c, d; Supplementary Movies M12-M14). These results indicate that DNMT1 and DOCK7 modulate mitochondrial positioning at nascent branch sites, which is negatively correlated with branch formation as reported by others^43^.

**Figure 7:**
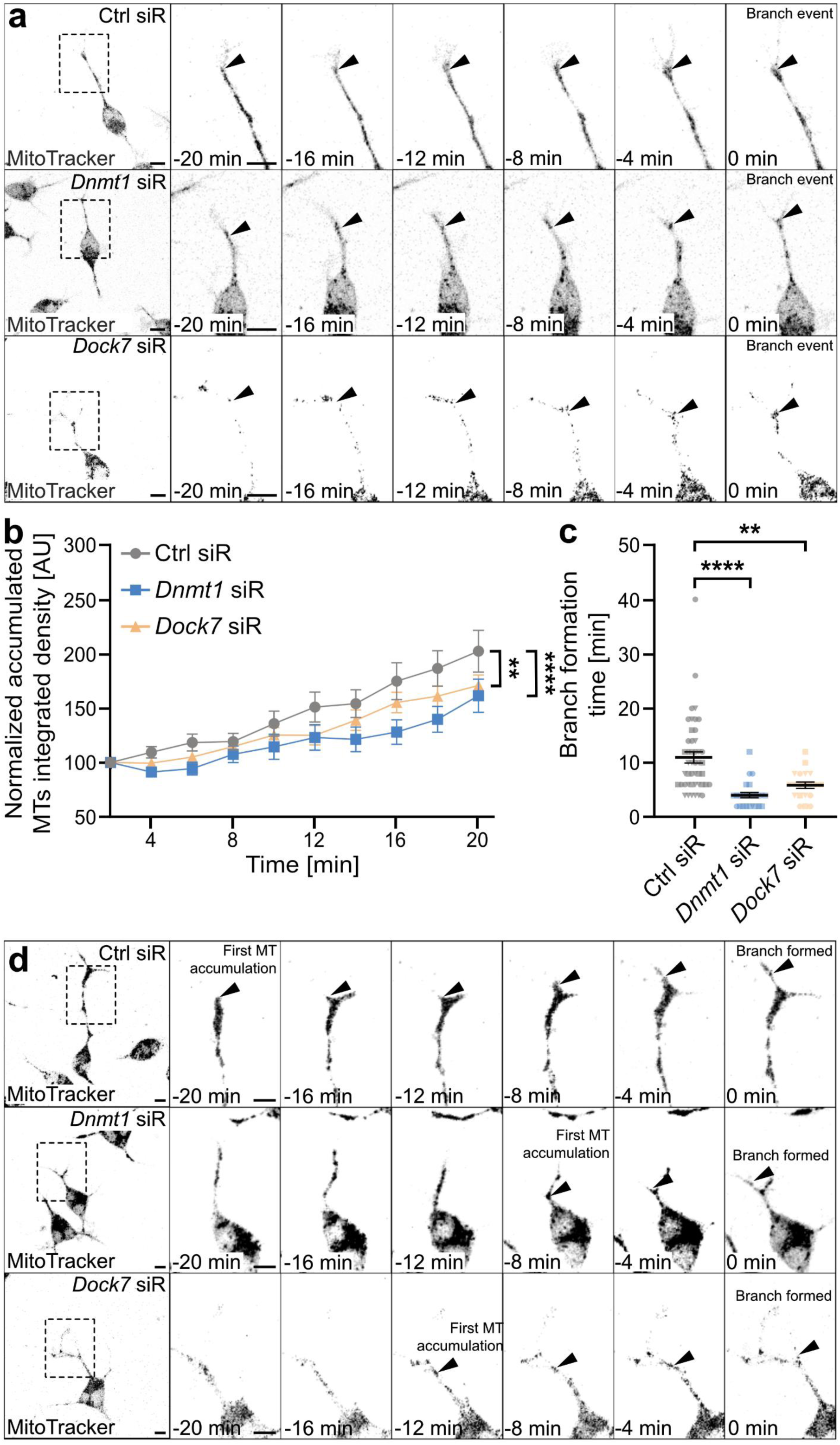
Mitochondria accumulation at prospective branching sites is impaired upon *Dnmt1* and *Dock7* depletion. (**a, b**) Live cell imaging analysis capturing mitochondria accumulation using the MitoTracker^TM^ Deep Red FM prior to branch formation in N2a cells 24 h after control, *Dnmt1*, or *Dock7* siRNA transfection. The white squares in the first panel indicate the magnified regions shown for each time frame. **(a)** Inverted grayscale images of mitochondria accumulation at branch initiation sites, Scale bars: 10 µm. The MitoTracker^TM^ integrated density, normalized to the corresponding integrated density of the same position in the first frame, at prospective branchpoints is shown in (**b**). n (Ctrl siR) = 22 cells with 54 events; n (*Dnmt1* siR) = 13 cells with 24 events; n (*Dock7* siR) = 21 cells with 43 events. N = 4 experiments. Two-way ANOVA followed by Tukey’s post-hoc multiple comparison test, ** p < 0.01. (**c, d**) Analysis of the branch formation time (**c**) and exemplary tracking of a branching event (from the first mitochondria accumulation puncta to the formation of the branch) (**d**). The white squares in the first panel indicate the magnified regions shown for each time frame. Scale bars: 20 µm. n (Ctrl siR) = 22 cells with 54 events; n (*Dnmt1* siR) = 13 cells with 24 events; n (*Dock7* siR) = 21 cells with 43 events. N = 4 experiments. One-way ANOVA followed by Dunnett’s post-hoc multiple comparison test, ** p < 0.01, **** p < 0.0001. Data are presented as mean ± SEM.

### DNMT1 and DOCK7 regulate microtubule stability, converging on STMN1 phosphorylation

Mitochondria and other organelles are trafficked along microtubules, whose post-translational modifications (PTMs) critically regulate transport. α-tubulin K40-acetylation provides stable tracks that enhance kinesin- and dynein-driven motility, whereas reduced acetylation disrupts trafficking. Reduced acetylation destabilizes the lattice, impairs efficient long-range trafficking and local retention, and at the same time increases microtubule dynamics, which has been linked to excessive or misregulated branching^44,45^.

We therefore analyzed α-tubulin acetylation levels in primary cortical neurons and N2a cells by immunocytochemistry. While total βIII-tubulin was unchanged, knockdown of both *Dnmt1* and *Dock7* significantly reduced acetylated α-tubulin in cortical neurons and N2a cells (Fig. 8a-d). This effect was not reproduced by pharmacological inhibition of DNA methylation using RG108 (Fig. 8e, f), supporting a DNA methylation-independent, likely cytosolic mechanism. Live imaging of EB3 comets in primary cortical neurons revealed no change in microtubule polymerization dynamics following knockdown of either *Dnmt1* or *Dock7* (Supplementary Fig. 7; Supplementary Movies M15-M17), indicating that neither protein affects microtubule growth per se.

**Figure 8:**
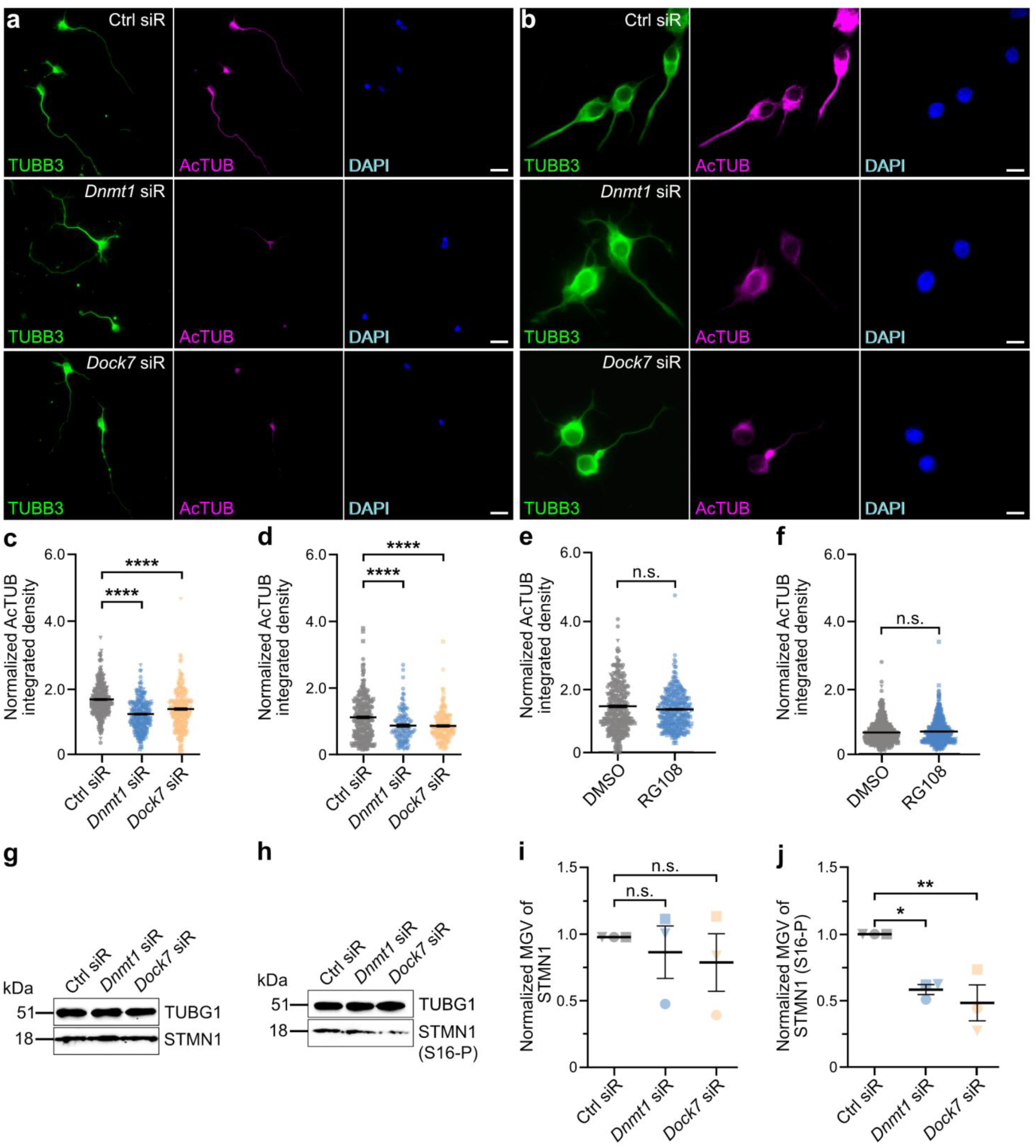
The acetylation of tubulin and phosphorylation of STMN1/Op18 (S16-P) are significantly reduced after depletion of *Dnmt1*/*Dock7* in cortical neurons (E14.5 + 2 DIV) and N2a cells. (**a, b**) Microphotographs of cortical neurons (**a,** Scale bars: 20 µm) and N2a cells (**b,** Scale bars: 40 µm), which were transfected with control, *Dnmt1*, or *Dock7* siRNA for 24 h after growing for one day, co-stained for DAPI (blue), ßIII-tubulin (TUBB3, green), and acetylated tubulin (AcTUB, magenta). (**c, d**) Analysis of the integrated density for the acetylated tubulin (AcTUB) normalized to the βIII-tubulin (TUBB3) integrated density after the respective knockdown in cortical neurons (**c**) and N2a cells (**d**). N = 3 experiments. One-way ANOVA followed by Dunnett’s post-hoc multiple comparison test, * p < 0.05, **** p < 0.0001. (**e, f**) Analysis of the acetylated tubulin (AcTUB) integrated density normalized to the βIII-tubulin (TUBB3) integrated density after the RG108 inhibitor treatment in cortical neurons (**e**) and N2a cells (**f**). N = 3 experiments. Two-tailed Student’s t-test, * p < 0.05. For (**c, e**): n (Ctrl siR) = 314 cells; n (*Dnmt1* siR) = 263 cells; n (*Dock7* siR) = 277 cells; n (DMSO) = 365 cells; n (RG108) = 358 cells. For (**d, f**): n (Ctrl siR) = 279 cells; n (*Dnmt1* siR) = 117 cells; n (*Dock7* siR) = 175 cells; n (DMSO) = 506 cells; n (RG108) = 525 cells. (**g, h**) Western blots revealing protein bands of STMN1 (**g**, TMW = 17 kDa) and the phosphorylated version of STMN1 (**h**, S16-P) (TMW = 17 kDa) in cortical single cell lysates (E14.5 + 2 DIV) treated previously with control, *Dnmt1*, or *Dock7* siRNA at 1 DIV for 24 h. γ-tubulin (**g**, **h**, TUBG1, TMW = 51 kDa) was used as a housekeeper. (**i, j**) Analysis of the mean grey value of STMN1 normalized against TUBG1 (**i**) and the mean grey value of the phosphorylated version of STMN1 (S16-P) (**j**) normalized against STMN1. N = 3 experiments. One-way ANOVA followed by Dunnett’s post-hoc multiple comparison test, * p < 0.05, ** p < 0.01. Data are presented as mean ± SEM.

Acetylation of α-tubulin is considered a PTM being associated with stable microtubules^46^. Given this relationship, the reduced acetylation signal seen after *Dnmt1* or *Dock7* knockdown points to less stable lattices. DOCK7 is known to stabilize microtubules by activating Rac/PAK signaling, which phosphorylates the microtubule-destabilizer STMN1 at Ser16 and thereby suppresses its activity^26^. Thus, we next asked whether DNMT1 also engages this pathway. Indeed, depletion of either *Dnmt1* or *Dock7* markedly reduced STMN1 phosphorylation (Ser16), while total STMN1 levels remained unchanged (Fig. 8g-j).

Our findings indicate that DNMT1 and DOCK7 converge on the same Rac/PAK–STMN1 axis. Under normal conditions, they promote STMN1 phosphorylation, thereby restraining its destabilizing activity and facilitating microtubule stability. Loss of either protein decreases STMN1 phosphorylation, which in turn reduces α-tubulin K40 acetylation, impairs organelle transport independently of cargo type, and promotes excessive branching.

Collectively, our data suggest that DNMT1 and DOCK7 form a cytosolic complex that converges on microtubule stability with implications for the regulation of branching in developing cortical neurons.

## Discussion

Neuronal morphogenesis relies on tightly regulated cytoskeletal remodeling, with microtubule dynamics shaping axons and dendrites^47^. While traditionally attributed to canonical cytosolic regulators, we identify an unexpected cytoplasmic role of the nuclear maintenance methyltransferase DNMT1 in coordinating microtubule stability and mitochondria trafficking, thereby constraining neurite branching in developing cortical neurons. DNMT1 acts independently of its methyltransferase activity by forming a complex with the polarity regulator DOCK7, converging on the Rac–STMN1 pathway to control mitochondrial transport. These findings redefine DNMT1 as a dual-compartment regulator that integrates structural and energetic modules during neuronal development.

Although classically known as the principal maintenance methyltransferase, DNMT1 possesses a large N-terminal domain rich in protein–protein interaction motifs^11,12,13^, consistent with scaffolding functions. Cytoplasmic DNMT1 has been observed in neurons and glia^19,20^, but its function has remained relatively unclear. Here, we demonstrate that catalytically inactive and cytosolic DNMT1 mutants fully rescue the branching phenotype caused by knockdown of endogenous *Dnmt1*, establishing extranuclear DNMT1 as essential for morphogenesis. This expands DNMT1’s role beyond transcriptional regulation^3,5,13^, placing it within a growing class of nuclear proteins repurposed for cytoskeletal signaling^1,48^.

Our proteomic and functional data identify DOCK7 as a key DNMT1 interactor. DOCK7 is known to regulate polarity and dendritic branching via Rac-dependent phosphorylation of STMN1, a microtubule-destabilizing protein^26,49^. In its unphosphorylated state, STMN1 destabilizes microtubules by sequestering soluble tubulin dimers and by increasing catastrophe rates^50,51^. Its phosphorylated form loses this ability, promoting stable microtubules^50,51^. Acetylation of α-tubulin is considered a PTM being associated with stable microtubules^46^. We show that depletion of either *Dnmt1* or *Dock7* enhances branching while reducing STMN1 phosphorylation and the levels of acetylated α-tubulin, without affecting polymerization dynamics. This suggests that DNMT1 and DOCK7 converge on the Rac– STMN1 pathway to maintain stable microtubules, and through this restrict excessive branching.

The link between α-tubulin acetylation and neuronal branching has been well-established: loss of α-tubulin acetylation leads to axon overbranching, which can be rescued by pharmacological stabilization of microtubules with taxol^52^. Thus, in our model, DNMT1 may serve as a scaffold anchoring DOCK7 to microtubules, enabling localized STMN1 phosphorylation and reinforcing microtubule stability. As no microtubule binding domain is described for DNMT1, this anchoring could occur via DNMT1 interaction with MAP2, a microtubule binding and stabilizing protein^53^, which we identified in our proteomic study. In this scenario, DNMT1 functions not as an enzymatic effector but as a structural adaptor, integrating structural and signaling modules to fine-tune cytoskeletal dynamics.

Mitochondrial positioning and trafficking are closely linked to the stability and post-translational modification of microtubules. Acetylated microtubules provide stable tracks that facilitate efficient organelle transport, whereas mitochondria–microtubule contacts can themselves promote α-tubulin acetylation^54,55^. This reciprocal relationship is coordinated by the outer-membrane GTPase MFN2, which recruits the acetyltransferase ATAT1 to mitochondria– microtubule contact sites, enhancing local α-tubulin acetylation and generating stabilized transport tracks^54^. Accordingly, the reduction in α-tubulin acetylation observed upon *Dnmt1* or *Dock7* depletion may reflect impaired microtubule stabilization and diminished mitochondria– microtubule coupling. Whether the trafficking phenotype arises primarily from microtubule destabilization or from altered mitochondria docking is beyond the scope of the current study. Nevertheless, our findings suggest that DNMT1 and DOCK7 jointly coordinate microtubule organization and mitochondrial deployment through a cytosolic scaffolding mechanism linking cytoskeletal stability and organelle positioning.

Future experimental efforts will have to address whether the branching phenotype is primarily driven by altered microtubule stability and resulting PTM^56^, or by altered mitochondria deployment^43^. Acetylation is known to protect microtubules from severing, while unacetylated lattices are more susceptible to katanin-mediated severing^57,58^. Thus, reduced acetylation following *Dnmt1* or *Dock7* depletion could increase severing activity, facilitating branch initiation. This is in line with recent work that emphasizes microtubule PTMs to govern neuronal branching. Loss of doublecortin promotes excessive branching through reduced tubulin polyglutamylation, impairing lysosomal trafficking^59^, while polymerization rates were not affected. Similarly, we find that *Dnmt1* or *Dock7* depletion increases branching despite unaltered EB3 dynamics, accompanied by decreased STMN1 phosphorylation, reduced tubulin acetylation, and altered organelle transport.

Apart from implications for neuronal development, the discovery of a cytosolic function of DNMT1 might also have profound implications for disease. DNMT1 mutations cause neurodegenerative syndromes such as Hereditary Sensory Neuropathy with Dementia and Hearing Loss (HSN-IE) and Autosomal Dominant Cerebellar Ataxia, Deafness, and Narcolepsy (ADCA-DN)^60,61^, conditions which are poorly explained by its canonical nuclear roles. Patient-derived fibroblasts carrying these mutations show mitochondrial hyper-function and elevated oxidative stress without altered mitochondrial DNA methylation^21^, suggesting the disruption of cytosolic DNMT1 functions. Importantly, a recent study revealed that DNMT1 also facilitates m⁵C RNA methylation by recruiting NSUN2 to mitochondrial metabolism-related transcripts, enhancing their stability^22^. Mutations in DNMT1’s RFTS domain cause aberrantly increased m⁵C RNA methylation and mitochondrial dysfunction^22^, shedding further light on how DNMT1’s extranuclear activities modulate mitochondrial health and contribute to neurodegeneration. Here, similar to our findings, DNMT1 function is mediated via its interaction with another protein, the RNA methyltransferase NSUN2. Additionally, the reported DNMT1 relocalization to the cytosol following brain injury^20^ may represent an adaptive mechanism aimed at stabilizing microtubules and preserving mitochondrial positioning under stress.

In summary, our findings uncover a DNA methylation-independent cytosolic role of DNMT1 in neuronal morphogenesis. By scaffolding DOCK7 and linking Rac–STMN1 signaling to microtubule stability and mitochondria trafficking, DNMT1 enforces structural and energetic checkpoints during branch formation. This work broadens the conceptual view of DNMT1 from a genome guardian to a dual-compartment regulator with key functions in cytoskeletal and organelle regulation, as well as broad implications for neurodevelopment and disease.

## Methods

### Structural predictions

The murine DOCK7/DNMT1 complex was modeled using a multi-step computational protocol. First, we refined the DNMT1 X-ray structure (residues 357-1612 (PDB ID: 3AV4^62^) and AlphaFold^63^ predicted DOCK7 model (residues 1-2130) in solution by explicit solvent molecular dynamics (MD) simulations (400 ns and 1000 ns for DNMT1 and DOCK7, respectively). 6 and 24 Na+ counterions were added to achieve neutrality of the system. The AMBER14SB^64^, TIP3P^65^, and Aqvist force fields^66^ were used for the proteins, water, and counter ions, respectively. Electrostatics were calculated using the Particle Mesh Ewald method^67^. A cutoff of 12 Å for the real part of electrostatics and for van der Waals was used. Energy minimization involved sequential steepest descent (5000 steps) followed by conjugate gradient (5000 steps) algorithms. Thermal equilibration involved heating from 0 K to 300 K over 5 ns under NVT ensemble conditions, followed by 50 ns of pressure equilibration under NPT ensemble using a Langevin thermostat^63^ (γ = 1.0 ps⁻¹) and a Berendsen barostat^68^ at 1 atm pressure. Then, 400 ns and 1000 ns production runs were carried out for DNMT1 and DOCK7, respectively. The k-means clustering of the trajectory in AMBER22^69^ generated 3 representative conformers for DNMT1 and 6 for DOCK7. Next, we used a multistep docking approach involving ClusPro^38^, LZerD^39^, RosettaDock^40^, and GRAMM^41^ to obtain 720 models of the complex. A hierarchical k-means clustering procedure identified 5 conformational families. The clustering centers were qualitatively energy-ranked by MM-GBSA binding free energy calculations. The structural determinants of the top two centers, each with a normalized affinity greater than 0.5, were investigated in this study. The hotspot interface residues for each model were determined through the MM-GBSA^69^ code, considering the top 10 polar residues ranked by energy contribution.

### Animals

Wildtype C57Bl/6J mice (Jackson, USA) were used for E14.5 cortical neuron and tissue dissections, maintained according to EU guidelines, FELASA, GV-SOLAS, and LANUV regulations. Mice were kept at 20-24 °C and 45-65% relative humidity. A day-night rhythm of 12 h was set. Food and water were provided *ad libitum*.

For mitochondrial and EB3 trafficking dynamics, neuronal morphology in P0 cortical neurons, and in utero electroporation, wildtype Jcl:ICR mice (Japan SLC, Japan) were used, maintained following the guidelines of the Animal Experiment Committee of the Kyoto University. Mice were kept at 23 ± 3 °C and 50% relative humidity. A day-night rhythm of 12 h was adjusted. Food and water were provided *ad libitum*.

### Isolation of murine E14.5 and P0 brains

E14.5 embryos were extracted as described in Reichard et al. (2025)^3^. A pregnant C57Bl/6J mouse was intraperitoneally injected with a lethal overdose of ketamine/xylazine and embryos were extracted via abdominal incision. The embryos were decapitated and cortices were collected in ice-cold 1x Geys Balanced Salt Solution (GBSS)/0.65% glucose.

P0 brains were extracted as described in Wu et al. (2015)^70^. Jcl:ICR pups were cryo-anesthetized and rapidly decapitated. Brains were extracted and collected in ice-cold 1x HBSS (Gibco, USA)/0.6% glucose.

### Single-cell dissociation of isolated E14.5 and P0 cortices

Extracted E14.5 cortices were dissociated as described in Reichard et al. (2025)^3^. Cortical single neurons were cultured in Neurobasal™ medium with 1x B-27™ and 0.25x GlutaMAX™ (Gibco, USA) and seeded at a density of 300 cells/mm² per laminin (19 μg/ml, Sigma Aldrich, USA) and poly-L-lysine (10 μg/ml, Sigma Aldrich, USA) coated coverslip. Transfection was performed at 1 DIV (for 2 DIV analyses) and 3 DIV (for analysis at 5 DIV).

P0 cortices were dissociated using the Neuron Dissociation kit (Fujifilm, Japan) following Hatsuda et al. (2023)^71^. Neurons were seeded with a density of 2655 cells/mm^2^ or 1695 cells/mm^2^ per coated glass base dish (IWAKI, Japan) for the mitochondrial trafficking or morphology studies, respectively, on laminin (19 μg/ml, Sigma Aldrich, USA) and poly-D-lysine (10 μg/ml, Sigma Aldrich, USA) coated surfaces. Cytarabine (AraC) was not used. Co-transfection occurs at 3 DIV (mitochondrial trafficking experiments) or 4 DIV (morphology studies). All cultures were maintained at 37 °C, 5% CO_2_, and 96% relative humidity.

### Cultivation of immortalized N2a cells

Immortalized neuroblastoma (N2a) cells (ATCC: CCL-131) were cultivated as described in Bayer et al. (2020)^72^ in Dulbecco’s modified Eagle’s medium (DMEM) (4.5 g/L D-glucose, GlutaMAX^TM^, pyruvate (Gibco, USA), 2% fetal bovine serum (FBS, Biowest, USA)) at 37 °C, 5% CO_2_, and 96% relative humidity. For thawing and maintenance cultures, penicillin (100 U/mL)/streptomycin (100 µg/mL) (P/S, Thermo Fisher Scientific, USA) was added. Cells were seeded at a density of 150 cells/mm² (RG108/CRISPRi-Ctrl/*Dnmt1*-GFP experiments), 110 cells/mm² (TMRE assay), and 75 cells/mm² (mitochondria accumulation assay) per laminin (19 μg/ml, Sigma Aldrich, USA) and poly-L-lysine (10 μg/ml, Sigma Aldrich, USA) coated coverslip or well.

### Transfection/Co-transfection of siRNA/plasmid DNA

Cells were transfected with *Dnmt1* (30 nM, Santa Cruz, USA, #sc-35203) or *Dock7* siRNA (30 nM, Santa Cruz, USA, #sc-105312) additionally with 15 nM of the control siRNA Block-it^TM^ control siRNA conjugated to Alexa555/488 (Thermo Fisher Scientific, USA, #14750100/#2013) or conjugated to Cy5 (Cell Signaling Technology, USA, #86921) using Lipofectamine^TM^ 2000 (Thermo Fisher Scientific, USA) following the manufacturer’s instructions. The following plasmid constructs were additionally co-transfected: MT-GFP (20 ng/µL), MT-dsRed (30 ng/µL), the respective CAG-mNG-*Dnmt1* expression construct (400 ng/µL), CAG-EB3-GFP (400 ng/µL), CAG-mSc (50 ng/µL), and CRISPRi-Ctrl/*Dnmt1*-GFP (400 ng/µL) (Supplementary Tab. 1).

### Treatment with RG108 and fixation

24 h after seeding, cells were treated with RG108 (25 µM, Abcam, USA, #ab141013) for 24 h. For immunocytochemical staining, cells were fixed with prewarmed (37 °C) 4% paraformaldehyde (PFA; Merck, USA) in 1x Phosphate-buffered saline (PBS) for 10 min.

### Treatment with TMRE and FCCP

For assay validation, cells were treated 24 h after seeding with either FCCP (2 µM, Abcam, USA, #ab120081) or DMSO for 40 min at 37 °C, followed by TMRE labeling (5 µM, Abcam, USA, #ab274305) for 20 min at 37 °C. The medium was replaced with fresh medium (w/o TMRE; w FCCP/DMSO). For the siRNA-mediated knockdown conditions, TMRE staining was performed 24 h post-transfection for 20 min, followed by medium change and imaging. Analysis was executed using *ImageJ* (NIH, USA) to analyze the TMRE integrated density, which was normalized to the mean of the control condition (background-corrected).

### Treatment with MitoTracker™ Deep Red FM

23 h after transfection, N2a cells were incubated with MitoTracker™ Deep Red FM (1:750; Thermo Fisher Scientific, USA, #M22426) for 30 min at 37 °C. The medium was replaced, and imaging was started. Mitochondrial accumulation was analyzed by measuring the integrated fluorescence densities at branch points (background corrected) using *ImageJ* (NIH, USA), comparing the onset of branch formation to the frame captured ten time points earlier. All integrated density values were normalized to the initial frame. Branch formation time was quantified in parallel.

### Plasmid constructs

The *Dnmt1* mutants were generated according to Zhou et al. (2024)^73^. *Dnmt1*s nuclear localization sequence and catalytic domain were identified using UniProt (UniProt, 2022) and NCBI databases. PCR-based insert amplification was performed using siRNA-resistant wildtype pENTR4-V5-*Dnmt1*-cDNA-for-n-Term-Fusion and pENTR4-V5-*Dnmt1*-ΔpCat (P1228G, C1229S) [15]. For the Dnmt1-NLS mutant, two fragments were synthesized independently and later ligated using the In-Fusion Cloning kit (Takara, Japan) into a pENTR1A backbone vector (Invitrogen, USA). The *Dnmt1* fragments were fused with the CAG-mNG destination vector using the Gateway^TM^ LR clonase^TM^ kit (Invitrogen, USA). For the *Dnmt1* repressive CRISPRi-Dnmt1-GFP construct, the gRNA sequence for DNMT1 (5′-ACCCCAGTAAACTAACCCCCGGG-3′) was cloned into the pX458-CAG-dCas9KRAB-2A-GFP vector as described in Hatsuda et al. (2023)^71^. All plasmids are listed in Supplementary Table 1.

### In utero electroporation and histology

The in utero electroporation was performed at E12.5 in pregnant Jcl:ICR mice as described by Kawaue et al. (2019)^74^. Plasmid DNA solution (5 µg/µl CRISPRi-Dnmt1/Ctrl-GFP), 0.2 µg/µl pCAG-mSc, and 0.01% Fast Green (Nacalai, Japan), were injected into the lateral ventricles using a pulled glass micropipette followed by 5 square pulses (32 V, 50 ms, 450 ms interval) via a CUY21 electroporator (NEPA Gene, Japan). Embryos were placed back into the abdominal cavity and the abdomen was sutured.

For immunohistochemistry, mice at P5 were perfused with PBS followed by 4% paraformaldehyde in phosphate buffer (PB). Their brains were removed and postfixed for 3-4 h at 4°C. After washing with PB, the brains were embedded in 3.5% low-melting agarose in PB. The brains were then sectioned into 300 μm-thick coronal slices with a vibratome (NLS-AT, Dosaka EM). The samples were cleared according to the product protocol (CUBIC Trial Kit, Fujifilm-WAKO, Japan). Briefly, the tissues were immersed in 1/2 CUBIC-1 solution (1:1 mixture of water and CUBIC-1) at RT for 20min, followed by CUBIC-1 solution with gentle shaking at 37°C for 15 min. The decolorized samples were washed three times with PB at RT under gentle shaking. The sections were blocked with 0.5% BSA and 0.1% Triton X-100 in PB at RT for 30 min. The sections were incubated with a chick anti-GFP (1:2000, Invitrogen, A10262) and anti-Ctip2(1:1000, BioLegend, 650601) or anti-DNMT1 primary antibody in blocking solution at 37°C for 1-2 days, followed by incubation with a goat anti-chicken Alexa Fluor 488 (1:500, Invitrogen, A11039) and a goat anti-rat Alexa Fluor 647 (1:500, Invitrogen, A21247) or a donkey anti-rabbit Alexa Fluor 647 (1:500, Invitrogen, A31573) secondary antibody at 37°C for 1-2 days. After washing with PB containing 0.1% Triton X-100, the samples were immersed in the 1/2 CUBIC-2 solution at RT for 5 min and then in the CUBIC-2 solution at 37°C for 30 min. Images of the fixed samples were acquired with a laser scanning confocal microscope (FV1000 and FV4000, Olympus) equipped with UPlanSApo 10× (NA 0.40) and UPLSAPO 40× (NA 0.95) objectives. For morphometric analysis, dendrites were traced using the Imaris software (Version 6.2.1, Oxford Instruments, UK) and the number of basal dendrites was counted manually.

### Immunocytochemical staining

Immunocytochemical staining without antigen retrieval was executed as described in Symmank et al. (2018)^10^. Primary antibodies (1x PBS/0.1% Triton X-100/4% BSA) were applied for 2 h at RT and secondary antibodies were applied for 1 h at RT. Nuclei were stained with 4’,6-diamidino-2-phenylindol dihydrochloride (DAPI, 1:10.000 in PBS; Carl Roth, Germany) and coverslips were embedded in Mowiol or Fluoromount-G™ (Thermo Fisher Scientific, USA, #495952). For morphological analysis of P0 + 5 DIV cortical neurons, DAPI staining was done directly after PFA fixation. Coverslips were washed with PBS/0.5% Triton X-100, stained with DAPI, rinsed and mounted with ProLong™ Gold (Thermo Fisher Scientific, USA, #P36934). Morphological parameters were manually analyzed using *ImageJ* (NIH, USA). The acetylated tubulin (AcTUB) integrated density was manually measured and normalized to the βIII-tubulin (TUBB3) integrated density using *ImageJ* (NIH, USA).

Antigen retrieval for 5mC and DNMT1 staining was performed as described in Pensold et al.(2021)^75^. Permeabilization (1x PBS/0.5% Triton X-100) and blocking (1x PBS/0.5% Triton X-100/4% BSA) were performed as described above. For quantification, the 5mC/DNMT1 integrated density was manually measured and normalized to the DAPI integrated density using *ImageJ* (NIH, USA). For the colocalization studies between DNMT1 and DOCK7, antibodies were conjugated according to the FlexAble CoraLite® Plus 647 (Proteintech, USA, #KFA003) and FlexAble CoraLite® Plus 555 (Proteintech, USA, #KFA002) kits following the manufacturer’s protocol (2.5 µg antibody each). Conjugated antibodies were incubated for 2 h at RT on permeabilized cells, followed by DAPI staining and mounting with ProLong™ Diamond (Thermo Fisher Scientific, USA, #P36965). Colocalization was analyzed following Costes threshold method [20] in *Imaris* (Oxford Instruments, UK). Specificity was achieved by comparing signal overlap in defined cytosolic volumes to a 90°-rotated control channel, revealing non-specific colocalization.

The following primary antibodies were used: mouse anti-5mC (1:250; Diagenode, Belgium, #C15200081), mouse anti-SMI-312 (1:500; Biolegend, USA, #837904), mouse anti-acetylated-tubulin (1:2000; Sigma Aldrich, USA, #T6793), rabbit anti-DNMT1 (1:1000; BioAcademia, Japan, #70-201), mouse anti-SERCA2 (1:1000; Santa Cruz, USA, #sc-376235), mouse anti-LAMP1 (1:1000; Santa Cruz, USA, #sc-20011), rabbit anti-DOCK7 (1:1000; Proteintech, USA, #1300-1-AP), chicken anti-GFP (1:500; Molecular Probes, USA, #A10262), and rabbit anti-mouse anti-βIII-tubulin (1:1000; Sigma Aldrich, USA, #T2200). The following secondary antibodies were used: Alexa488-goat anti-rabbit IgG (1:500; Thermo Fisher Scientific, USA, #A11008), Cy5-goat anti-mouse IgG (1:500; Jackson, USA, #115175146), and Cy5-goat anti-rabbit IgG (1:1000; Thermo Fisher Scientific, USA, #A10523).

Detailed information about the microscopes and the settings for capturing immunocyto- and histochemical stainings and live cell imaging are provided in the supplemental information.

### Sample collection and lysis

For the analysis of phosphorylated STMN1 after siRNA-mediated knockdown of *Dnmt1*/*Dock7*, E14.5 + 2 DIV cortical neurons were washed with prewarmed 1x DPBS (Merck, USA), mechanically detached, collected and pooled in maintenance medium, and centrifuged (800 x g, 4 min, 4 °C), followed by a DPBS washing and snap-freezing. For DNMT1/DOCK7 co-immunoprecipitation, E14.5 cortical tissue was dissected in 1x HBSS/0.65% glucose on ice and processed similarly.

Co-immunoprecipitated samples were treated with lysis buffer (0.1 M Na_2_HPO_4_, 0.1 M NaH_2_PO_4_, 5 mM EDTA, 2 mM MgCl_2_, 0.1% CHAPS) containing 1% leupeptin (1 mg/ml, Serva, Germany) and 1% Phenylmethylsulfonyl Fluoride (PMSF, 100 mM, Sigma Aldrich, USA). For STMN1 phosphorylation analysis, Radio immunoprecipitation (RIPA) buffer (150 mM NaCl, 50 mM Tris-HCl, 1% NP-40, 0.5% CHAPS, 0.1% SDS, 1 mM EDTA, 10 mM NaF), including leupeptin and PMSF was used. Samples were homogenized for 1 h on ice using a douncing homogenizer (Fisherbrand, USA) and then centrifuged for 20 min at 13.000 x g at 4 °C.

### Subcellular fractionation

Cytoplasmic and nuclear fractions were prepared based on the protocol by Suzuki et al. (2010)^76^. N2a cells (∼95% confluency) were washed, detached in cold 1x PBS, and centrifuged. The cell pellets were resuspended in 1 ml of 0.1% NP40/PBS (Whole-cell lysate, WCL). After centrifugation, the supernatant was collected (cytoplasmic fraction). The remaining pellet was resuspended in 0.1% NP40/PBS and divided into two samples (nuclear fraction): One pellet was resuspended in 1x SDS sample buffer; the other pellet was resuspended 0.1% NP40/PBS. All lysates and nuclear samples were sonicated (10 × 30 s), after which the SDS samples were denaturated. Non-SDS samples were centrifuged (8.000 x g, 30 s, 4 °C), and the immunoprecipitation was continued.

### Co-Immunoprecipitation (Co-IP)

Protein A-agarose beads (Merck, USA) were washed twice with Co-IP buffer (20 mM HEPES, 0.1 mM EDTA, 50 mM KCl, 0.05% CHAPS), then incubated and mixed with 3 µg rabbit anti-DNMT1 or rabbit IgG in 10 µl Co-IP buffer for 1 h on ice and blocked with 45 µg BSA for 1 h. After washing, 1 mg of total protein, measured using the Qubit™ Protein Assay Kit and Qubit™ 4 fluorometer (Thermo Fisher Scientific, USA), was added per sample and incubated overnight at 4 °C on a rotator. Beads were then washed six times with Co-IP buffer. For mass spectrometry, 5 mg (brain tissue, 3.5 months) or 3.65 mg (N2a cells) of total protein was used and prepared as described above.

### Protein sample denaturation, SDS-PAGE, and Western blot

For DNMT1/IgG immunoprecipitates, 4x SDS sample buffer (200 mM Tris-HCl, 4% SDS, 10% β-mercaptoethanol, 40% glycerol, 0.002% bromophenol blue) was added, and samples were denatured at 95 °C for 10 min. For STMN1 assays, 1 mg of protein input was used. Proteins were separated using the Gel™ HSE or Gel™ TG PRiME™ 4-20% gradient gels (Serva, Germany). The following SDS-PAGE (Laemmli buffer, Serva, Germany) and protein band transfer via semi-dry blotting (Towbin blotting kit, Serva,Germany) were performed according to the manufacturer’s protocol. Membranes were blocked and incubated with antibodies as described in Bayer et al. (2020)^72^. For reprobing, membranes were stripped (2 × 7 min, mild stripping buffer (1.5% Glycine, 0.1% SDS, 1% Tween® 20; pH 2.2)), washed in PBS (2 × 10 min) and TBS-T (2 × 5 min) at RT and then re-incubated. Protein band quantification was performed using *Image Lab* (6.0.1, Bio-Rad Laboratories, USA) and *ImageJ* (1.52p, NIH, USA). STMN1 levels were normalized to γ-tubulin (TUBG1), and phosphorylated STMN1 (S16-P) was normalized to total STMN1.

The following primary antibodies were used: rabbit anti-DNMT1 (1:1000; BioAcademia, Japan, #70-201), rabbit anti-DOCK7 (1:1000; Proteintech, USA, #1300-1-AP), rabbit anti-IgG (1:500; Merck, USA, #12-370), rabbit anti-phosphorylated STMN1/Op18 (1:500; Abcam, USA, #ab47328), rabbit anti-STMN1/Op18 (1:500; Thermo Fisher Scientific, USA, #PA5-28092), and mouse anti-γ-tubulin (1:1000; Sigma Aldrich, USA, #T6557). The following secondary antibodies were used: HRP-sheep anti-mouse (1:4000; Cytiva, USA, #NA931) and HRP-donkey anti-rabbit (1:4000; Cytiva, USA, #NA9340).

### Mass spectrometry

For mass spectrometry, DNMT1 co-immunoprecipitated proteins were separated by SDS-PAGE and visualized using Quick Coomassie Stain (Serva, Germany) according to the manufacturer’s protocol. Sample preparation and analysis were performed by the IZKF Core Facility (University Hospital RWTH Aachen, Germany) as described in Heymann et al. (2019)^77^. Measurements were conducted using a mass spectrometer (Q Exactive Hybrid Quadrupole-Orbitrap, Thermo Fisher Scientific, USA). Data analysis was executed using Perseus software (Max-Planck-Institute of Biochemistry, Germany). The Gene Ontology (GO) enrichment analysis was performed using STRING and *ShinyGO 0.82*^78,79^.

### Statistics and figure illustration

Statistical analyses were performed using *GraphPad Prism* (9.5.1, GraphPad Software, USA). For normally distributed data, two-tailed Student’s t-test or Welch’s t-test for comparisons between two groups, two-way ANOVA followed by Tukey’s post hoc multiple comparisons test, or one-way ANOVA followed by Dunnett’s post hoc test (for comparisons involving three groups) or Tukey’s post hoc test (for more than three groups) were applied. For non-normally distributed data, the Kruskal-Wallis test followed by Dunn’s post hoc test was executed. Significance levels, number of replicates (N), and the number of analyzed cells (n) are provided in the figure legends for each graph. All experiments were independently repeated at least three times. The central line of the scatter dot blots represents the median, including the standard error of the mean (±SEM). Symbols were used to represent data sets from individual experiments. Figures were illustrated using *Affinity* (2.6, Serif, Great Britain).

### Data Availability

The data supporting the results of this study are available as described in the results and the method part or were directly uploaded to corresponding data platforms. We confirm there is no (privacy) conflict of sharing our data openly. Since our study did not used human/patient material there is no need to anonymize corresponding data to comply with ethical and legal standards. For further information regarding data access or requests, please contact Geraldine Zimmer-Bensch. The predicted structural determinants of the DNMT1/DOCK7 complex have been deposited in the Zenodo repository: https://doi.org/10.5281/zenodo.17342500.

### Code Availability

No in house code was developed for this research. All the codes used are freely available for academic research.

## Acknowledgements

This work was supported by the Proteomics and the Flow Cytometry Facility, two core facilities of the Interdisciplinary Center for Clinical Research (IZKF) Aachen within the Faculty of Medicine at RWTH Aachen University (DFG project number: 439895892).

We thank Sandra Brill for her support with mouse care.

This research was funded by the Deutsche Forschungsgemeinschaft (DFG, German Research Foundation)—368482240/GRK2416 dedicated to G.Z.B and M.S., as well as ZI-1224/13-1, ZI-1224/19-1 dedicated to G.Z.B; the study was further funded under the Excellence Strategy of the Federal Government and the Länder (OPSF678, OPSF812, SFASIA002) dedicated to G.Z.B.; K.Z. received financial support from the European Union−NextGenerationEU−Project Title PpiAsd (Protein−protein Interactions in Autism Spectrum Disorder)−CUP F53D23001170006. K.Z. and P.R. acknowledge the support from HEAL ITALIA (project number PE00000019) and eINS Ecosystem of Innovation for Next Generation Sardinia (CUP: F53C22000430001) within the framework of the National Recovery and Resilience Plan (NRRP).

## Author contributions Statement

G.P. performed experiments and data analysis, figure illustration, assisted in manuscript writing and editing; J.K. and T.K. performed in vivo experiments, imaging and data analysis; K.Z. performed in silico experiments, data analysis, and manuscript editing; S.X. performed in silico experiments, data analysis, and figure illustration, and manuscript editing; C.W.H. contributed to data acquisition and analysis; C.B. K. performed experiments and data analysis; J.D. performed experiments and data analysis, figure illustration; P.W. performed experiments and data analysis, figure illustration; S.N performed in vivo experiments, imaging and data analysis; C.P.S. performed experiments and data analysis; A.R. performed experiments and data analysis, figure illustration; J.E.W. performed experiments and data analysis, figure illustration; P.R. Performed data analysis, and manuscript editing; M.S. contributed to data interpretation, supervision, and manuscript editing; P.C. performed conceptual design of in silico experiments, and wrote the in silico part of the manuscript; M.K., contributed to data interpretation, supervision, and manuscript editing. G.Z.B. conducted the overall conceptual design, administration and supervision, experimental wet-lab design, wrote the manuscript, figure editing, data analysis, interpretation and discussion.

