## Supplementary Figures, Legends, information and methods for "A cytosolic function of DNMT1 controls neuronal morphogenesis via microtubule regulation"

#### 1 Supplementary Figure Legends

**Supplementary Figure S1: Knockdown of *Dnmt1* using the CRISPRi-Dnmt1-GFP plasmid increases branching of cortical neurons (E14.5 + 2 DIV) and N2a cells in vitro.** (**a, b**) Images of N2a cells immunocytochemically stained for DNMT1 (magenta) and DAPI (blue), 24 h after the transfection with either the CRISPRi-Ctrl-GFP (**a**) or CRISPRi-Dnmt1-GFP (**b**) plasmid (green). Successfully transfected N2a cells are marked with white arrowheads. Scale bars: 5 $\mu$m. (**c**) Quantification of the DNMT1 integrated density normalized to the DAPI integrated density. n (CRISPRi-Ctrl-GFP) = 144 cells; n (CRISPRi-Dnmt1-GFP) = 150 cells. N = 3 experiments. (**d, e**) Inverted microphotographs of βIII-tubulin immunocytochemically stained cortical neurons (E14.5 + 2 DIV), transfected with CRISPRi-Ctrl-GFP (**d**) or CRISPRi-Dnmt1-GFP (**e**) plasmids at 1 DIV for 24 h Scale bars: 20 $\mu$m. (**f-h**) Analysis of the length of the longest processes (**f**), the total number of all primary processes (**g**), and the branches normalized to the sum of all process lengths (**h**). n (CRISPRi-Ctrl-GFP) = 64 cells; n (CRISPRi-Dnmt1-GFP) = 66 cells. N = 3 experiments. (**i, j**) Inverted microphotographs of βIII-tubulin stained N2a cells transfected with either CRISPRi-Ctrl-GFP (**i**) or CRISPRi-Dnmt1-GFP (**j**) for 24 h. Scale bars: 20 $\mu$m. (**k-m**) Analysis of the length of the longest process (**k**), the total number of primary processes (**l**), and the branches normalized to the sum of all process lengths (**m**). n (CRISPRi-Ctrl-GFP) = 106 cells; n (CRISPRi-Dnmt1-GFP) = 96 cells. N = 3 experiments. (**n**) Immunocytochemical staining for βIII-tubulin (TUBB3, green), SMI-312 (red), and DAPI (blue). Scale bar: 10 $\mu$m. (**o**) Analysis of how many of the longest processes of cortical neurons (E14.5 + 2 DIV) were SMI-312-positive and presumably axons. n = 95 cells. N = 3 independent experiments. Two-tailed Student's t-test, * $p < 0.05$, **** $p < 0.0001$. Data are presented as mean ± SEM.

**Supplementary Figure S2: Knockdown of *Dnmt1* using siRNA increases branching of cortical neurons (P0 + 5 DIV) in vitro.** (**a, b**) Inverted microphotographs of exemplary βIII-tubulin immunohistochemically stained cortical neurons (E14.5 + 5 DIV) transfected with control (**a**) or *Dnmt1* (**b**) siRNA, 48 h after transfection at DIV 3 for further morphological analysis. Scale bars: 20 $\mu$m. (**c-e**) Analysis of the length of the longest process (**c**), the number of processes (**d**), and the branches normalized to the longest process length likely representing axon (**e**). n (Ctrl siR) = 33 cells; n (*Dnmt1* siR) = 45 cells. N = 3 experiments (**f, g**) Inverted microphotographs of exemplary CAG-mSc-co-transfected cortical neurons (P0 + 5 DIV) with control (**f**) or *Dnmt1* (**g**) siRNA, 24 h after transfection at DIV 4 for further morphological analysis. Scale bars: 50 $\mu$m. (**h-j**) Analysis of the length of the longest process (**h**), the number of processes (**i**), and the branches normalized to the longest process length likely representing axon (**j**). n (Ctrl siR) = 48 cells; n (*Dnmt1* siR) = 73 cells. N = 3 experiments. Two-tailed Student's t-test, * $p < 0.05$, *** $p < 0.001$. Data are presented as mean ± SEM.

**Supplementary Figure S3: The DNMT inhibitor RG108 successfully blocks DNA methylation in N2a cells and does not impair the neuronal morphology of cortical neurons at E14.5 + 2 DIV and DNMT1 is localized in the cytosol.** (**a-c**) Validation of the DNMT inhibitor RG108. N2a cells were treated 24 h after seeding either with DMSO, as a control, or RG108 for 24 h. Afterwards, immunocytochemical staining using a specific antibody targeting 5mC and DAPI staining were executed. In (**a, b**), exemplary thermal images of the

5mC fluorescence signal are shown for the DMSO (**a**) and RG108 (**b**) conditions in N2a cells. The corresponding analysis of 5mC integrated densities normalized to the DAPI integrated densities of N2a cells is shown in (**c**). n (DMSO) = 155 cells; n (RG108) = 155 cells. N = 3 experiments. (**d, e**) Inverted exemplary microphotographs of DMSO (**d**) or RG108 (**e**) treated cortical neurons (E14.5 + 2 DIV) 24 h after the respective treatments at 1 DIV and immunocytochemically stained for βIII-tubulin after 2 DIV. Scale bars: 20 $\mu$m. (**f-h**) Analysis of morphological parameters, including the length of the longest process (**f**), the total number of processes (**g**), and the number of all branches normalized to the lengths summed up over all processes (**h**). n (DMSO) = 167 cells; n (RG108) = 164 cells. N = 3 experiments. Two-tailed Student's t-test, * $p < 0.05$, **** $p < 0.0001$. Data are presented as mean ± SEM. (**i**) Protein detection of DNMT1, ACTB, and H2AX in N2a cells by Western blot following immunoprecipitation using either a DNMT1-specific antibody or an IgG control antibody, after subcellular fractionation.

**Supplementary Figure S4: DNMT1 interacts with proteins related to intracellular trafficking and mitochondria.** (**a, b**) List of DNMT1 interacting cytosolic proteins, which show a fold change of at least two in murine brain tissue (3.5 months, Supplementary data 1) and N2a cells (Supplementary data 2), related to intracellular trafficking (**a**) and mitochondria (**b**). (**c**) Venn diagram representing the overlap between proteins being co-immunoprecipitated with DNMT1 in N2a cell lysates (Supplementary data 2) and human cell lysates as described in Wang et al., 2025. (**d, e**) Co-immunocytochemically staining of N2a cells growing for 48 h with DAPI (blue) and antibodies against DNMT1 (red) and the endoplasmic reticulum (SERCA2, green) (**d**) or lysosomes (LAMP1, green) (**e**) using high-resolution STED microscopy for colocalization analysis. Scale bars: 5 $\mu$m. The white squares are depicting the magnification of the merge the merge. Scale bars: 2 $\mu$m.

**Supplementary Figure S5: Depletion of *Dnmt1* does not impair the mitochondria size or TMRE signal**. (**a-c**) Quantitative analysis of the general mitochondria lengths (**a**), including both anterograde (**b**) and retrograde (**c**) transported mitochondria and their respective lengths in cortical neurons (P0 + 4 DIV). n (Ctrl siR) = 15 cells with 128 MTs; n (*Dnmt1* siR) = 11 cells with 147 MTs. N = 3 experiments. One-way ANOVA followed by Dunnett's post-hoc multiple comparison test, * $p < 0.05$. (**d-f**) Validation of the TMRE labeling. (**d, e**) Exemplary thermal images of the TMRE signal after treating N2a cells 48 h after seeding with DMSO as a control (**d**) or the ATPase inhibitor FCCP (**e**). The quantification of the TMRE integrated density normalized to the mean of the DMSO control after the treatments of N2a cells is shown in (**f**). n (DMSO) = 149 cells; n (FCCP) = 149 cells. N = 3 experiments. Two-tailed Student's t-test, **** $p < 0.0001$. (**g**) Representative microphotographs of N2a cells treated with TMRE (red) one day after control or *Dnmt1* siRNA transfection. Scale bars: 25 $\mu$m. (**h**) Quantitative analysis of the TMRE integrated density normalized to the mean of the control group in N2a cells transfected after one day either with control or *Dnmt1* siRNA for 24 h. n (Ctrl siR) = 400 cells; n (*Dnmt1* siR) = 400 cells. N = 4 experiments. Kruskal-Wallis test, * $p < 0.05$. Data are presented as mean ± SEM.

**Supplementary Figure S6: Depletion of *Dock7* do not affect the mitochondria length or the TMRE signal, but elevates the trafficking velocity of endolysosomal organelles.** (**a**, **b**) Inverted microphotographs of exemplary βIII-tubulin stained N2a cells transfected with either control (**a**) or *Dock7* siRNA (**b**) for 24 h. Scale bars: 20 µm. (**c-e**) Analysis of the length of the longest process (**c**), the total number of processes (**d**), and the branches normalized to the sum of all process lengths (**e**). n (Ctrl siR) = 166 cells; n (*Dock7* siR) = 166 cells. N = 3 experiments. (**f-h**) Quantitative analysis of the general mitochondria lengths (**f**), including both anterograde (**g**) and retrograde (**h**) transported mitochondria and their respective lengths in cortical neurons (P0 + 4 DIV). n (Ctrl siR) = 15 cells with 128 MTs; n (*Dock7* siR) = 11 cells with 142 MTs). N = 3 experiments. Two-tailed Student's t-test, * $p < 0.05$. (**i**) Representative microphotographs of N2a cells treated with TMRE (red) one day after control or *Dock7* siRNA transfection. Scale bars: 25 µm. (**j**) Analysis of the TMRE integrated density normalized to the mean of the control group in N2a cells transfected after one day either with control or *Dock7* siRNA for 24 h. n (Ctrl siR) = 400 cells, n (*Dock7* siR) = 400 cells. N = 4 experiments. Kruskal-Wallis test, * $p < 0.05$. Data are presented as mean ± SEM.

**Supplementary Figure S7: Depletion of *Dnmt1* or *Dock7* do not affect the EB3 comet dynamics.** (**a**) Exemplary images of a co-transfected cortical neuron (P0 + 4 DIV) with Alexa Fluor 555 labeled control siRNA (red) and the EB3-mNG plasmid (green) 24 h after co-transfection at DIV 3. Scale bar: 5 µm. The white square represents the magnification of the merge. Scale bar: 2.5 µm. (**b**) Exemplary kymographs of EB3 comet trafficking after the knockdown of *Dnmt1* or *Dock7* are shown. (**c, d**) Quantitative analysis of the EB3 comet velocity (**c**) and the relative proportion of EB3 comets' distal moving direction (**d**). n (Ctrl siR) = 12 cells with 213 EB3 comets; n (*Dnmt1* siR) = 12 cells with 218 EB3 comets, n (*Dock7* siR) = 10 cells with 181 EB3 comets. N = 3 experiments. One-way ANOVA followed by Dunnett's post-hoc multiple comparison test, * $p < 0.05$.

#### 2 Supplementary Movie Legend

**Supplementary Movie M1:** Representative live cell imaging video of a cortical neuron (P0 + 4 DIV) co-transfected with control siRNA and MT-dsRed at 3 DIV for 24 h. Scale bar: 5 µm.

**Supplementary Movie M2:** Representative live cell imaging video of a cortical neuron (P0 + 4 DIV) co-transfected with *Dnmt1* siRNA and MT-dsRed at 3 DIV for 24 h. Scale bar: 5 µm.

**Supplementary Movie M3:** Representative live cell imaging video of a cortical neuron (P0 + 4 DIV) co-transfected with *Dnmt1* siRNA, MT-dsRed, and the Dnmt1-WT expression construct at 3 DIV for 24 h. Scale bar: 5 µm.

**Supplementary Movie M4:** Representative live cell imaging video of a cortical neuron (P0 + 4 DIV) co-transfected with *Dnmt1* siRNA, MT-dsRed, and the Dnmt1-ΔpCat mutant construct at 3 DIV for 24 h. Scale bar: 5 µm.

**Supplementary Movie M5:** Representative live cell imaging video of a cortical neuron (P0 + 4 DIV) co-transfected with *Dnmt1* siRNA, MT-dsRed, and the Dnmt1-ΔNLS mutant construct at 3 DIV for 24 h. Scale bar: 5 µm.

**Supplementary Movie M6:** Representative live cell imaging video of a cortical neuron (P0 + 4 DIV) co-transfected with control siRNA and MT-GFP at 3 DIV for 24 h. Scale bar: 5 µm.

**Supplementary Movie M7:** Representative live cell imaging video of a cortical neuron (P0 + 4 DIV) co-transfected with *Dnmt1* siRNA and MT-GFP at 3 DIV for 24 h. Scale bar: 5 $\mu$m.

**Supplementary Movie M8:** Representative live cell imaging video of a cortical neuron (P0 + 4 DIV) co-transfected with *Dock7* siRNA and MT-GFP at 3 DIV for 24 h. Scale bar: 5 $\mu$m.

**Supplementary Movie M9:** Representative live cell imaging video (inverted) of a N2a cell (grown for 48 h) transfected with control siRNA and treated with *MitoTracker™ Deep Red FM* 24 h after transfection. Scale bar: 10 $\mu$m.

**Supplementary Movie M10:** Representative live cell imaging video (inverted) of a N2a cell (grown for 48 h) transfected with *Dnmt1* siRNA and treated with *MitoTracker™ Deep Red FM* 24 h after transfection. Scale bar: 10 $\mu$m.

**Supplementary Movie M11:** Representative live cell imaging video (inverted) of a N2a cell (grown for 48 h) transfected with *Dock7* siRNA and treated with *MitoTracker™ Deep Red FM* 24 h after transfection. Scale bar: 10 $\mu$m.

**Supplementary Movie M12:** Representative live cell imaging video (inverted) of a N2a cell (grown for 48 h) transfected with control siRNA and treated with *MitoTracker™ Deep Red FM* 24 h after transfection for the analysis of the branch formation time. Scale bar: 10 $\mu$m.

**Supplementary Movie M13:** Representative live cell imaging video (inverted) of a N2a cell (grown for 48 h) transfected with *Dnmt1* siRNA and treated with *MitoTracker™ Deep Red FM* 24 h after transfection for the analysis of the branch formation time. Scale bar: 10 $\mu$m.

**Supplementary Movie M14:** Representative live cell imaging video (inverted) of a N2a cell (grown for 48 h) transfected with *Dock7* siRNA and treated with *MitoTracker™ Deep Red FM* 24 h after transfection for the analysis of the branch formation time. Scale bar: 10 $\mu$m.

**Supplementary Movie M15:** Representative live cell imaging video of a cortical neuron (P0 + 4 DIV) co-transfected with control siRNA and EB3-GFP at 3 DIV for 24 h. Scale bar: 5 $\mu$m.

**Supplementary Movie M16:** Representative live cell imaging video of a cortical neuron (P0 + 4 DIV) co-transfected with *Dnmt1* siRNA and EB3-GFP at 3 DIV for 24 h. Scale bar: 5 $\mu$m.

**Supplementary Movie M17:** Representative live cell imaging video of a cortical neuron (P0 + 4 DIV) co-transfected with *Dock7* siRNA and EB3-GFP at 3 DIV for 24 h. Scale bar: 5 $\mu$m.

#### 3 Supplementary Tables

**Supplementary table 1:**
**All used plasmid constructs in this paper and their corresponding origins.**

| Plasmid name | Origin |
|---|---|
| pAAV-CAG-Mito-dsRed (MT-dsRed) | Fukumitsu et al. [16] |
| pAAV-CAG-Mito-EGFP (MT-GFP) | Fukumitsu et al. [16] |
| pCAG-mNG-Dest. | Zhou et al. [14] |
| pCAG-GFP-EB3 | Wu et al. [17] |
| pCAG-mNG-V5-Dnmt1-WT | This paper |
| pCAG-mNG-V5-Dnmt1-ΔCat (Δ1-1437) | This paper |
| pCAG-mNG-V5-Dnmt1-ΔNLS (Δ525-610) | This paper |

| | |
|---|---|
| pCAG-mNG-V5-Dnmt1-ΔpCat (P1228G, C1229S) | This paper |
| pCAG-mScarlet | Gift from Mineko Kengaku (Kyoto University, Japan) |
| pENTR4-V5-Dnmt1-cDNA-for-n-Term-Fusion (WT) | Haggerty et al. [15] |
| pENTR4-V5-Dnmt1-ΔpCat (P1228G, C1229S) | Haggerty et al. [15] |
| pX458-CAG-dCas9KRAB-2A-Dnmt1-GFP (CRISPRi-Dnmt1-GFP) | This paper |
| pX458-CAG-dCas9KRAB-2A-GFP-Empty (CRISPRi-Ctrl-GFP) | Hatsuda et al. [12] |
| CD63-pEGFP C2 | Gift from Paul Luzio (Addgene plasmid # 62964; http://n2t.net/addgene:62964; RRID: Addgene_62964) |
| pLAMP1-mCherry | Gift from Amy Palmer (Addgene plasmid #45147; http://n2t.net/addgene:45147; RRID: Addgene_45147) |

**Supplementary table 2:**

The *Dnmt1* cDNA sequence alignment here is shown for Dnmt1-WT/ΔNLS (Δ525-610)/ΔpCat (P1228G, C1229S)/ΔCat (Δ3426-4863) and was performed using the Multiple Sequence Alignment (MSA) online platform Clustal Omega[1]. Deletions and sequence changes (mutations) are highlighted in yellow.

```
Dnmt1-WT      GGATCACCTGCAAGGACAGCTCCAGCAAGAGTGCCCGCACTTGCCTCTCCTGCCGGAAGC    60
Dnmt1-ΔCat    GGATCACCTGCAAGGACAGCTCCAGCAAGAGTGCCCGCACTTGCCTCTCCTGCCGGAAGC    60
Dnmt1-ΔpCat   GGATCACCTGCAAGGACAGCTCCAGCAAGAGTGCCCGCACTTGCCTCTCCTGCCGGAAGC    60
Dnmt1-ΔNLS    GGATCACCTGCAAGGACAGCTCCAGCAAGAGTGCCCGCACTTGCCTCTCCTGCCGGAAGC    60

Dnmt1-WT      CTGCCTGACCATGTTAGGCGACGATTGAAAGACTTGGAGCGCGATGGACTCACCGAGAAA    120
Dnmt1-ΔCat    CTGCCTGACCATGTTAGGCGACGATTGAAAGACTTGGAGCGCGATGGACTCACCGAGAAA    120
Dnmt1-ΔpCat   CTGCCTGACCATGTTAGGCGACGATTGAAAGACTTGGAGCGCGATGGACTCACCGAGAAA    120
Dnmt1-ΔNLS    CTGCCTGACCATGTTAGGCGACGATTGAAAGACTTGGAGCGCGATGGACTCACCGAGAAA    120

Dnmt1-WT      GAATGCGTGAGAGAGAAACTGAACCTTTTGCATGAATTCCTTCAAACCGAAATTAAGAGC    180
Dnmt1-ΔCat    GAATGCGTGAGAGAGAAACTGAACCTTTTGCATGAATTCCTTCAAACCGAAATTAAGAGC    180
Dnmt1-ΔpCat   GAATGCGTGAGAGAGAAACTGAACCTTTTGCATGAATTCCTTCAAACCGAAATTAAGAGC    180
Dnmt1-ΔNLS    GAATGCGTGAGAGAGAAACTGAACCTTTTGCATGAATTCCTTCAAACCGAAATTAAGAGC    180

Dnmt1-WT      CAATTGTGCGATTTGGAAACCAAGCTGCATAAGGAAGAACTTTCCGAAGAGGGCTATTTG    240
Dnmt1-ΔCat    CAATTGTGCGATTTGGAAACCAAGCTGCATAAGGAAGAACTTTCCGAAGAGGGCTATTTG    240
Dnmt1-ΔpCat   CAATTGTGCGATTTGGAAACCAAGCTGCATAAGGAAGAACTTTCCGAAGAGGGCTATTTG    240
Dnmt1-ΔNLS    CAATTGTGCGATTTGGAAACCAAGCTGCATAAGGAAGAACTTTCCGAAGAGGGCTATTTG    240

Dnmt1-WT      GCCAAAGTAAAATCCTTGCTGAACAAAGATTTGTCACTGGAGAATGGCACCCACACATTG    300
Dnmt1-ΔCat    GCCAAAGTAAAATCCTTGCTGAACAAAGATTTGTCACTGGAGAATGGCACCCACACATTG    300
Dnmt1-ΔpCat   GCCAAAGTAAAATCCTTGCTGAACAAAGATTTGTCACTGGAGAATGGCACCCACACATTG    300
Dnmt1-ΔNLS    GCCAAAGTAAAATCCTTGCTGAACAAAGATTTGTCACTGGAGAATGGCACCCACACATTG    300

Dnmt1-WT      ACTCAAAAAGCCAATGGTTGTCCCGCAAACGGCTCTCGACCCACCTGGCGCGCCGAAATG    360
Dnmt1-ΔCat    ACTCAAAAAGCCAATGGTTGTCCCGCAAACGGCTCTCGACCCACCTGGCGCGCCGAAATG    360
Dnmt1-ΔpCat   ACTCAAAAAGCCAATGGTTGTCCCGCAAACGGCTCTCGACCCACCTGGCGCGCCGAAATG    360
Dnmt1-ΔNLS    ACTCAAAAAGCCAATGGTTGTCCCGCAAACGGCTCTCGACCCACCTGGCGCGCCGAAATG    360

Dnmt1-WT      GCTGACTCTAATAGATCCCCACGCTCAAGACCAAAACCAAGAGGCCCTAGACGATCCAAG    420
Dnmt1-ΔCat    GCTGACTCTAATAGATCCCCACGCTCAAGACCAAAACCAAGAGGCCCTAGACGATCCAAG    420
Dnmt1-ΔpCat   GCTGACTCTAATAGATCCCCACGCTCAAGACCAAAACCAAGAGGCCCTAGACGATCCAAG    420
Dnmt1-ΔNLS    GCTGACTCTAATAGATCCCCACGCTCAAGACCAAAACCAAGAGGCCCTAGACGATCCAAG    420

Dnmt1-WT      AGTGATTCTGATACACTGTCAGTTGAAACCTCCCCATCATCCGTGGCTACAAGGCGAACC    480
Dnmt1-ΔCat    AGTGATTCTGATACACTGTCAGTTGAAACCTCCCCATCATCCGTGGCTACAAGGCGAACC    480
Dnmt1-ΔpCat   AGTGATTCTGATACACTGTCAGTTGAAACCTCCCCATCATCCGTGGCTACAAGGCGAACC    480
Dnmt1-ΔNLS    AGTGATTCTGATACACTGTCAGTTGAAACCTCCCCATCATCCGTGGCTACAAGGCGAACC    480

Dnmt1-WT      ACCAGGCAAACAACAATTACTGCACACTTCACTAAAGGTCCTACCAAACGGAAACCTAAG    540
```

```
Dnmt1-ΔCat    ACCAGGCAAACAACAATTACTGCACACTTCACTAAAGGTCCTACCAAACGGAAACCTAAG    540
Dnmt1-ΔpCat   ACCAGGCAAACAACAATTACTGCACACTTCACTAAAGGTCCTACCAAACGGAAACCTAAG    540
Dnmt1-ΔNLS    ACCAGGCAAACAACAATTACTGCACACTTCACTAAAGGTCCTACCG--------------    526

Dnmt1-WT      GAGGAGAGTGAAGAAGGTAATTCCGCCGAGAGTGCAGCAGAGGAGCGGGACCAAGACAAG    600
Dnmt1-ΔCat    GAGGAGAGTGAAGAAGGTAATTCCGCCGAGAGTGCAGCAGAGGAGCGGGACCAAGACAAG    600
Dnmt1-ΔpCat   GAGGAGAGTGAAGAAGGTAATTCCGCCGAGAGTGCAGCAGAGGAGCGGGACCAAGACAAG    600
Dnmt1-ΔNLS    ------------------------------------------------------------    526

Dnmt1-WT      AAGAGGAGGGTAGTCGACACCGAAAGCGGAGCCGCTGCCGCAGTAGAAAAGCTTGAAGAA    660
Dnmt1-ΔCat    AAGAGGAGGGTAGTCGACACCGAAAGCGGAGCCGCTGCCGCAGTAGAAAAGCTTGAAGAA    660
Dnmt1-ΔpCat   AAGAGGAGGGTAGTCGACACCGAAAGCGGAGCCGCTGCCGCAGTAGAAAAGCTTGAAGAA    660
Dnmt1-ΔNLS    ----------TAGTCGACACCGAAAGCGGAGCCGCTGCCGCAGTAGAAAAGCTTGAAGAA    576

Dnmt1-WT      GTAACAGCTGGTACACAACTTGGGCCTGAAGAACCCTGTGAACAAGAGGATGACAACCGG    720
Dnmt1-ΔCat    GTAACAGCTGGTACACAACTTGGGCCTGAAGAACCCTGTGAACAAGAGGATGACAACCGG    720
Dnmt1-ΔpCat   GTAACAGCTGGTACACAACTTGGGCCTGAAGAACCCTGTGAACAAGAGGATGACAACCGG    720
Dnmt1-ΔNLS    GTAACAGCTGGTACACAACTTGGGCCTGAAGAACCCTGTGAACAAGAGGATGACAACCGG    636

Dnmt1-WT      AGTCTGAGGCGCCACACCCGCGAGCTGAGCCTGAGAAGGAAATCCAAAGAAGACCCTGAT    780
Dnmt1-ΔCat    AGTCTGAGGCGCCACACCCGCGAGCTGAGCCTGAGAAGGAAATCCAAAGAAGACCCTGAT    780
Dnmt1-ΔpCat   AGTCTGAGGCGCCACACCCGCGAGCTGAGCCTGAGAAGGAAATCCAAAGAAGACCCTGAT    780
Dnmt1-ΔNLS    AGTCTGAGGCGCCACACCCGCGAGCTGAGCCTGAGAAGGAAATCCAAAGAAGACCCTGAT    696

Dnmt1-WT      CGGGAGGCTCGGCCCGAGACTCACCTGGATGAGGACGAAGACGGGAAAAAGGATAAACGG    840
Dnmt1-ΔCat    CGGGAGGCTCGGCCCGAGACTCACCTGGATGAGGACGAAGACGGGAAAAAGGATAAACGG    840
Dnmt1-ΔpCat   CGGGAGGCTCGGCCCGAGACTCACCTGGATGAGGACGAAGACGGGAAAAAGGATAAACGG    840
Dnmt1-ΔNLS    CGGGAGGCTCGGCCCGAGACTCACCTGGATGAGGACGAAGACGGGAAAAAGGATAAACGG    756

Dnmt1-WT      AGCAGTCGGCCACGGTCTCAGCCACGCGATCCCGCAGCAAAACGACGGCCCAAAGAGGCA    900
Dnmt1-ΔCat    AGCAGTCGGCCACGGTCTCAGCCACGCGATCCCGCAGCAAAACGACGGCCCAAAGAGGCA    900
Dnmt1-ΔpCat   AGCAGTCGGCCACGGTCTCAGCCACGCGATCCCGCAGCAAAACGACGGCCCAAAGAGGCA    900
Dnmt1-ΔNLS    AGCAGTCGGCCACGGTCTCAGCCACGCGATCCCGCAGCAAAACGACGGCCCAAAGAGGCA    816

Dnmt1-WT      GAGCCCGAGCAGGTGGCACCCGAGACTCCCGAAGATCGCGATGAGGATGAACGCGAAGAG    960
Dnmt1-ΔCat    GAGCCCGAGCAGGTGGCACCCGAGACTCCCGAAGATCGCGATGAGGATGAACGCGAAGAG    960
Dnmt1-ΔpCat   GAGCCCGAGCAGGTGGCACCCGAGACTCCCGAAGATCGCGATGAGGATGAACGCGAAGAG    960
Dnmt1-ΔNLS    GAGCCCGAGCAGGTGGCACCCGAGACTCCCGAAGATCGCGATGAGGATGAACGCGAAGAG    876

Dnmt1-WT      AAGCGACGGAAGACCACACGCAAAAAGCTCGAGTCCCATACTGTGCCCGTTCAAAGTCGA    1020
Dnmt1-ΔCat    AAGCGACGGAAGACCACACGCAAAAAGCTCGAGTCCCATACTGTGCCCGTTCAAAGTCGA    1020
Dnmt1-ΔpCat   AAGCGACGGAAGACCACACGCAAAAAGCTCGAGTCCCATACTGTGCCCGTTCAAAGTCGA    1020
Dnmt1-ΔNLS    AAGCGACGGAAGACCACACGCAAAAAGCTCGAGTCCCATACTGTGCCCGTTCAAAGTCGA    936

Dnmt1-WT      TCAGAGCGAAAGGCCGCACAGTCAAAGAGCGTTATACCTAAGATAAACTCACCTAAGTGT    1080
Dnmt1-ΔCat    TCAGAGCGAAAGGCCGCACAGTCAAAGAGCGTTATACCTAAGATAAACTCACCTAAGTGT    1080
Dnmt1-ΔpCat   TCAGAGCGAAAGGCCGCACAGTCAAAGAGCGTTATACCTAAGATAAACTCACCTAAGTGT    1080
Dnmt1-ΔNLS    TCAGAGCGAAAGGCCGCACAGTCAAAGAGCGTTATACCTAAGATAAACTCACCTAAGTGT    996

Dnmt1-WT      CCTGAATGTGGGCAGCACCTTGATGATCCAAACCTTAAATATCAGCAACATCCTGAAGAC    1140
Dnmt1-ΔCat    CCTGAATGTGGGCAGCACCTTGATGATCCAAACCTTAAATATCAGCAACATCCTGAAGAC    1140
Dnmt1-ΔpCat   CCTGAATGTGGGCAGCACCTTGATGATCCAAACCTTAAATATCAGCAACATCCTGAAGAC    1140
Dnmt1-ΔNLS    CCTGAATGTGGGCAGCACCTTGATGATCCAAACCTTAAATATCAGCAACATCCTGAAGAC    1056

Dnmt1-WT      GCCGTTGATGAGCCACAAATGCTTACTTCAGAAAAATTGTCAATATACGATTCCACTTCA    1200
Dnmt1-ΔCat    GCCGTTGATGAGCCACAAATGCTTACTTCAGAAAAATTGTCAATATACGATTCCACTTCA    1200
Dnmt1-ΔpCat   GCCGTTGATGAGCCACAAATGCTTACTTCAGAAAAATTGTCAATATACGATTCCACTTCA    1200
Dnmt1-ΔNLS    GCCGTTGATGAGCCACAAATGCTTACTTCAGAAAAATTGTCAATATACGATTCCACTTCA    1116

Dnmt1-WT      ACTTGGTTCGACACTTATGAAGACTCACCTATGCATAGGTTTACATCCTTTTCCGTATAT    1260
Dnmt1-ΔCat    ACTTGGTTCGACACTTATGAAGACTCACCTATGCATAGGTTTACATCCTTTTCCGTATAT    1260
Dnmt1-ΔpCat   ACTTGGTTCGACACTTATGAAGACTCACCTATGCATAGGTTTACATCCTTTTCCGTATAT    1260
Dnmt1-ΔNLS    ACTTGGTTCGACACTTATGAAGACTCACCTATGCATAGGTTTACATCCTTTTCCGTATAT    1176

Dnmt1-WT      TGCAGCAGAGGTCACCTCTGCCCAGTGGATACCGGGCTCATAGAGAAGAACGTCGAGCTG    1320
Dnmt1-ΔCat    TGCAGCAGAGGTCACCTCTGCCCAGTGGATACCGGGCTCATAGAGAAGAACGTCGAGCTG    1320
Dnmt1-ΔpCat   TGCAGCAGAGGTCACCTCTGCCCAGTGGATACCGGGCTCATAGAGAAGAACGTCGAGCTG    1320
Dnmt1-ΔNLS    TGCAGCAGAGGTCACCTCTGCCCAGTGGATACCGGGCTCATAGAGAAGAACGTCGAGCTG    1236

Dnmt1-WT      TATTTCTCAGGTTGCGCAAAGGCAATTCATGACGAGAACCCTTCAATGGAAGGAGGCATT    1380
Dnmt1-ΔCat    TATTTCTCAGGTTGCGCAAAGGCAATTCATGACGAGAACCCTTCAATGGAAGGAGGCATT    1380
Dnmt1-ΔpCat   TATTTCTCAGGTTGCGCAAAGGCAATTCATGACGAGAACCCTTCAATGGAAGGAGGCATT    1380
Dnmt1-ΔNLS    TATTTCTCAGGTTGCGCAAAGGCAATTCATGACGAGAACCCTTCAATGGAAGGAGGCATT    1296

Dnmt1-WT      AACGGTAAAAACCTTGGTCCCATTAACCAGTGGTGGCTTAGTGGCTTTGACGGAGGAGAG    1440
Dnmt1-ΔCat    AACGGTAAAAACCTTGGTCCCATTAACCAGTGGTGGCTTAGTGGCTTTGACGGAGGAGAG    1440
Dnmt1-ΔpCat   AACGGTAAAAACCTTGGTCCCATTAACCAGTGGTGGCTTAGTGGCTTTGACGGAGGAGAG    1440
```

```
Dnmt1-ΔNLS    AACGGTAAAAACCTTGGTCCCATTAACCAGTGGTGGCTTAGTGGCTTTGACGGAGGAGAG    1356

Dnmt1-WT      AAGGTCCTCATTGGGTTTAGCACCGCCTTTGCCGAATACATACTTATGGAGCCATCAAAG    1500
Dnmt1-ΔCat    AAGGTCCTCATTGGGTTTAGCACCGCCTTTGCCGAATACATACTTATGGAGCCATCAAAG    1500
Dnmt1-ΔpCat   AAGGTCCTCATTGGGTTTAGCACCGCCTTTGCCGAATACATACTTATGGAGCCATCAAAG    1500
Dnmt1-ΔNLS    AAGGTCCTCATTGGGTTTAGCACCGCCTTTGCCGAATACATACTTATGGAGCCATCAAAG    1416

Dnmt1-WT      GAGTACGAGCCTATTTTTGGACTTATGCAGGAAAAAATCTATATTAGCAAGATTGTCGTG    1560
Dnmt1-ΔCat    GAGTACGAGCCTATTTTTGGACTTATGCAGGAAAAAATCTATATTAGCAAGATTGTCGTG    1560
Dnmt1-ΔpCat   GAGTACGAGCCTATTTTTGGACTTATGCAGGAAAAAATCTATATTAGCAAGATTGTCGTG    1560
Dnmt1-ΔNLS    GAGTACGAGCCTATTTTTGGACTTATGCAGGAAAAAATCTATATTAGCAAGATTGTCGTG    1476

Dnmt1-WT      GAATTCCTCCAAAATAACCCCGACGCAGTGTATGAGGACCTGATAAATAAAATCGAGACT    1620
Dnmt1-ΔCat    GAATTCCTCCAAAATAACCCCGACGCAGTGTATGAGGACCTGATAAATAAAATCGAGACT    1620
Dnmt1-ΔpCat   GAATTCCTCCAAAATAACCCCGACGCAGTGTATGAGGACCTGATAAATAAAATCGAGACT    1620
Dnmt1-ΔNLS    GAATTCCTCCAAAATAACCCCGACGCAGTGTATGAGGACCTGATAAATAAAATCGAGACT    1536

Dnmt1-WT      ACAGTGCCACCATCAACTATAAACGTCAATCGGTTTACAGAAGACTCATTGTTGAGGCAT    1680
Dnmt1-ΔCat    ACAGTGCCACCATCAACTATAAACGTCAATCGGTTTACAGAAGACTCATTGTTGAGGCAT    1680
Dnmt1-ΔpCat   ACAGTGCCACCATCAACTATAAACGTCAATCGGTTTACAGAAGACTCATTGTTGAGGCAT    1680
Dnmt1-ΔNLS    ACAGTGCCACCATCAACTATAAACGTCAATCGGTTTACAGAAGACTCATTGTTGAGGCAT    1596

Dnmt1-WT      GCCCAGTTTGTGGTTTCCCAGGTAGAATCTTACGACGAGGCTAAAGATGATGACGAAACC    1740
Dnmt1-ΔCat    GCCCAGTTTGTGGTTTCCCAGGTAGAATCTTACGACGAGGCTAAAGATGATGACGAAACC    1740
Dnmt1-ΔpCat   GCCCAGTTTGTGGTTTCCCAGGTAGAATCTTACGACGAGGCTAAAGATGATGACGAAACC    1740
Dnmt1-ΔNLS    GCCCAGTTTGTGGTTTCCCAGGTAGAATCTTACGACGAGGCTAAAGATGATGACGAAACC    1656

Dnmt1-WT      CCTATCTTTCTGTCTCCATGTATGCGCGCACTGATCCATCTCGCTGGCGTGAGTCTCGGG    1800
Dnmt1-ΔCat    CCTATCTTTCTGTCTCCATGTATGCGCGCACTGATCCATCTCGCTGGCGTGAGTCTCGGG    1800
Dnmt1-ΔpCat   CCTATCTTTCTGTCTCCATGTATGCGCGCACTGATCCATCTCGCTGGCGTGAGTCTCGGG    1800
Dnmt1-ΔNLS    CCTATCTTTCTGTCTCCATGTATGCGCGCACTGATCCATCTCGCTGGCGTGAGTCTCGGG    1716

Dnmt1-WT      CAAAGAAGGGCAACAAGGCGGGTAATGGGGGCCACAAAAGAGAAGGACAAGGCACCCACT    1860
Dnmt1-ΔCat    CAAAGAAGGGCAACAAGGCGGGTAATGGGGGCCACAAAAGAGAAGGACAAGGCACCCACT    1860
Dnmt1-ΔpCat   CAAAGAAGGGCAACAAGGCGGGTAATGGGGGCCACAAAAGAGAAGGACAAGGCACCCACT    1860
Dnmt1-ΔNLS    CAAAGAAGGGCAACAAGGCGGGTAATGGGGGCCACAAAAGAGAAGGACAAGGCACCCACT    1776

Dnmt1-WT      AAAGCTACCACCACCAAACTCGTATATCAAATTTTCGATACCTTTTTCAGTGAACAAATC    1920
Dnmt1-ΔCat    AAAGCTACCACCACCAAACTCGTATATCAAATTTTCGATACCTTTTTCAGTGAACAAATC    1920
Dnmt1-ΔpCat   AAAGCTACCACCACCAAACTCGTATATCAAATTTTCGATACCTTTTTCAGTGAACAAATC    1920
Dnmt1-ΔNLS    AAAGCTACCACCACCAAACTCGTATATCAAATTTTCGATACCTTTTTCAGTGAACAAATC    1836

Dnmt1-WT      GAGAAATATGATAAAGAAGACAAAGAAAATGCTATGAAGCGGAGACGATGTGGAGTTTGT    1980
Dnmt1-ΔCat    GAGAAATATGATAAAGAAGACAAAGAAAATGCTATGAAGCGGAGACGATGTGGAGTTTGT    1980
Dnmt1-ΔpCat   GAGAAATATGATAAAGAAGACAAAGAAAATGCTATGAAGCGGAGACGATGTGGAGTTTGT    1980
Dnmt1-ΔNLS    GAGAAATATGATAAAGAAGACAAAGAAAATGCTATGAAGCGGAGACGATGTGGAGTTTGT    1896

Dnmt1-WT      GAAGTTTGTCAGCAGCCTGAGTGTGGAAAGTGCAAGGCCTGCAAAGATATGGTCAAGTTC    2040
Dnmt1-ΔCat    GAAGTTTGTCAGCAGCCTGAGTGTGGAAAGTGCAAGGCCTGCAAAGATATGGTCAAGTTC    2040
Dnmt1-ΔpCat   GAAGTTTGTCAGCAGCCTGAGTGTGGAAAGTGCAAGGCCTGCAAAGATATGGTCAAGTTC    2040
Dnmt1-ΔNLS    GAAGTTTGTCAGCAGCCTGAGTGTGGAAAGTGCAAGGCCTGCAAAGATATGGTCAAGTTC    1956

Dnmt1-WT      GGGGGTACTGGAAGAAGCAAACAGGCCTGTCTCAAGCGGCGATGCCCAAATCTCGCTGTC    2100
Dnmt1-ΔCat    GGGGGTACTGGAAGAAGCAAACAGGCCTGTCTCAAGCGGCGATGCCCAAATCTCGCTGTC    2100
Dnmt1-ΔpCat   GGGGGTACTGGAAGAAGCAAACAGGCCTGTCTCAAGCGGCGATGCCCAAATCTCGCTGTC    2100
Dnmt1-ΔNLS    GGGGGTACTGGAAGAAGCAAACAGGCCTGTCTCAAGCGGCGATGCCCAAATCTCGCTGTC    2016

Dnmt1-WT      AAGGAAGCCGATGACGACGAGGAAGCAGACGACGACGTAAGCGAGATGCCCAGTCCTAAG    2160
Dnmt1-ΔCat    AAGGAAGCCGATGACGACGAGGAAGCAGACGACGACGTAAGCGAGATGCCCAGTCCTAAG    2160
Dnmt1-ΔpCat   AAGGAAGCCGATGACGACGAGGAAGCAGACGACGACGTAAGCGAGATGCCCAGTCCTAAG    2160
Dnmt1-ΔNLS    AAGGAAGCCGATGACGACGAGGAAGCAGACGACGACGTAAGCGAGATGCCCAGTCCTAAG    2076

Dnmt1-WT      AAGCTGCATCAAGGCAAAAAGAAGAAGCAAAATAAAGATCGAATCTCATGGCTCGGCCAA    2220
Dnmt1-ΔCat    AAGCTGCATCAAGGCAAAAAGAAGAAGCAAAATAAAGATCGAATCTCATGGCTCGGCCAA    2220
Dnmt1-ΔpCat   AAGCTGCATCAAGGCAAAAAGAAGAAGCAAAATAAAGATCGAATCTCATGGCTCGGCCAA    2220
Dnmt1-ΔNLS    AAGCTGCATCAAGGCAAAAAGAAGAAGCAAAATAAAGATCGAATCTCATGGCTCGGCCAA    2136

Dnmt1-WT      CCAATGAAGATAGAAGAAAACAGGACTTACTACCAGAAGGTCAGCATCGACGAAGAAATG    2280
Dnmt1-ΔCat    CCAATGAAGATAGAAGAAAACAGGACTTACTACCAGAAGGTCAGCATCGACGAAGAAATG    2280
Dnmt1-ΔpCat   CCAATGAAGATAGAAGAAAACAGGACTTACTACCAGAAGGTCAGCATCGACGAAGAAATG    2280
Dnmt1-ΔNLS    CCAATGAAGATAGAAGAAAACAGGACTTACTACCAGAAGGTCAGCATCGACGAAGAAATG    2196

Dnmt1-WT      CTGGAAGTCGGCGATTGTGTTAGCGTGATCCCCGACGATTCCTCCAAGCCATTGTATCTG    2340
Dnmt1-ΔCat    CTGGAAGTCGGCGATTGTGTTAGCGTGATCCCCGACGATTCCTCCAAGCCATTGTATCTG    2340
Dnmt1-ΔpCat   CTGGAAGTCGGCGATTGTGTTAGCGTGATCCCCGACGATTCCTCCAAGCCATTGTATCTG    2340
Dnmt1-ΔNLS    CTGGAAGTCGGCGATTGTGTTAGCGTGATCCCCGACGATTCCTCCAAGCCATTGTATCTG    2256
```

| | | |
|---|---|---|
| Dnmt1-WT | GCTCGCGTAACAGCTTTGTGGGAAGATAAAAACGGACAGATGATGTTCCATGCCCACTGG | 2400 |
| Dnmt1-ΔCat | GCTCGCGTAACAGCTTTGTGGGAAGATAAAAACGGACAGATGATGTTCCATGCCCACTGG | 2400 |
| Dnmt1-ΔpCat | GCTCGCGTAACAGCTTTGTGGGAAGATAAAAACGGACAGATGATGTTCCATGCCCACTGG | 2400 |
| Dnmt1-ΔNLS | GCTCGCGTAACAGCTTTGTGGGAAGATAAAAACGGACAGATGATGTTCCATGCCCACTGG | 2316 |
| Dnmt1-WT | TTCTGTGCAGGAACCGACACCGTACTGGGAGCCACAAGTGATCCCCTGGAGCTCTTCCTG | 2460 |
| Dnmt1-ΔCat | TTCTGTGCAGGAACCGACACCGTACTGGGAGCCACAAGTGATCCCCTGGAGCTCTTCCTG | 2460 |
| Dnmt1-ΔpCat | TTCTGTGCAGGAACCGACACCGTACTGGGAGCCACAAGTGATCCCCTGGAGCTCTTCCTG | 2460 |
| Dnmt1-ΔNLS | TTCTGTGCAGGAACCGACACCGTACTGGGAGCCACAAGTGATCCCCTGGAGCTCTTCCTG | 2376 |
| Dnmt1-WT | GTGGGCGAATGTGAGAATATGCAGCTGTCTTACATACACTCAAAGGTCAAAGTTATATAC | 2520 |
| Dnmt1-ΔCat | GTGGGCGAATGTGAGAATATGCAGCTGTCTTACATACACTCAAAGGTCAAAGTTATATAC | 2520 |
| Dnmt1-ΔpCat | GTGGGCGAATGTGAGAATATGCAGCTGTCTTACATACACTCAAAGGTCAAAGTTATATAC | 2520 |
| Dnmt1-ΔNLS | GTGGGCGAATGTGAGAATATGCAGCTGTCTTACATACACTCAAAGGTCAAAGTTATATAC | 2436 |
| Dnmt1-WT | AAGGCACCCTCTGAGAATTGGGCCATGGAGGGTGGAACTGACCCTGAGACTACCCTCCCC | 2580 |
| Dnmt1-ΔCat | AAGGCACCCTCTGAGAATTGGGCCATGGAGGGTGGAACTGACCCTGAGACTACCCTCCCC | 2580 |
| Dnmt1-ΔpCat | AAGGCACCCTCTGAGAATTGGGCCATGGAGGGTGGAACTGACCCTGAGACTACCCTCCCC | 2580 |
| Dnmt1-ΔNLS | AAGGCACCCTCTGAGAATTGGGCCATGGAGGGTGGAACTGACCCTGAGACTACCCTCCCC | 2496 |
| Dnmt1-WT | GGAGCAGAAGATGGAAAAACCTATTTCTTCCAGCTCTGGTATAATCAGGAATACGCCAGG | 2640 |
| Dnmt1-ΔCat | GGAGCAGAAGATGGAAAAACCTATTTCTTCCAGCTCTGGTATAATCAGGAATACGCCAGG | 2640 |
| Dnmt1-ΔpCat | GGAGCAGAAGATGGAAAAACCTATTTCTTCCAGCTCTGGTATAATCAGGAATACGCCAGG | 2640 |
| Dnmt1-ΔNLS | GGAGCAGAAGATGGAAAAACCTATTTCTTCCAGCTCTGGTATAATCAGGAATACGCCAGG | 2556 |
| Dnmt1-WT | TTCGAAAGTCCACCCAAAACTCAGCCAACCGAGGATAACAAGCACAAATTTTGTCTGTCA | 2700 |
| Dnmt1-ΔCat | TTCGAAAGTCCACCCAAAACTCAGCCAACCGAGGATAACAAGCACAAATTTTGTCTGTCA | 2700 |
| Dnmt1-ΔpCat | TTCGAAAGTCCACCCAAAACTCAGCCAACCGAGGATAACAAGCACAAATTTTGTCTGTCA | 2700 |
| Dnmt1-ΔNLS | TTCGAAAGTCCACCCAAAACTCAGCCAACCGAGGATAACAAGCACAAATTTTGTCTGTCA | 2616 |
| Dnmt1-WT | TGTATACGATTGGCCGAACTTAGACAGAAAGAAATGCCAAAAGTCTTGGAACAGATCGAA | 2760 |
| Dnmt1-ΔCat | TGTATACGATTGGCCGAACTTAGACAGAAAGAAATGCCAAAAGTCTTGGAACAGATCGAA | 2760 |
| Dnmt1-ΔpCat | TGTATACGATTGGCCGAACTTAGACAGAAAGAAATGCCAAAAGTCTTGGAACAGATCGAA | 2760 |
| Dnmt1-ΔNLS | TGTATACGATTGGCCGAACTTAGACAGAAAGAAATGCCAAAAGTCTTGGAACAGATCGAA | 2676 |
| Dnmt1-WT | GAAGTGGACGGCCGCGTTTACTGCAGTAGCATTACAAAGAACGGTGTGGTTTACCGCCTT | 2820 |
| Dnmt1-ΔCat | GAAGTGGACGGCCGCGTTTACTGCAGTAGCATTACAAAGAACGGTGTGGTTTACCGCCTT | 2820 |
| Dnmt1-ΔpCat | GAAGTGGACGGCCGCGTTTACTGCAGTAGCATTACAAAGAACGGTGTGGTTTACCGCCTT | 2820 |
| Dnmt1-ΔNLS | GAAGTGGACGGCCGCGTTTACTGCAGTAGCATTACAAAGAACGGTGTGGTTTACCGCCTT | 2736 |
| Dnmt1-WT | GGTGATAGTGTATATCTTCCACCTGAAGCTTTTACTTTTAATATCAAGGTTGCAAGCCCC | 2880 |
| Dnmt1-ΔCat | GGTGATAGTGTATATCTTCCACCTGAAGCTTTTACTTTTAATATCAAGGTTGCAAGCCCC | 2880 |
| Dnmt1-ΔpCat | GGTGATAGTGTATATCTTCCACCTGAAGCTTTTACTTTTAATATCAAGGTTGCAAGCCCC | 2880 |
| Dnmt1-ΔNLS | GGTGATAGTGTATATCTTCCACCTGAAGCTTTTACTTTTAATATCAAGGTTGCAAGCCCC | 2796 |
| Dnmt1-WT | GTCAAGAGGCCTAAGAAGGACCCCGTGAACGAGACTCTCTACCCCGAACACTACCGGAAA | 2940 |
| Dnmt1-ΔCat | GTCAAGAGGCCTAAGAAGGACCCCGTGAACGAGACTCTCTACCCCGAACACTACCGGAAA | 2940 |
| Dnmt1-ΔpCat | GTCAAGAGGCCTAAGAAGGACCCCGTGAACGAGACTCTCTACCCCGAACACTACCGGAAA | 2940 |
| Dnmt1-ΔNLS | GTCAAGAGGCCTAAGAAGGACCCCGTGAACGAGACTCTCTACCCCGAACACTACCGGAAA | 2856 |
| Dnmt1-WT | TATTCAGACTACATTAAAGGGAGTAACCTCGACGCACCTGAGCCCTATCGGATTGGACGG | 3000 |
| Dnmt1-ΔCat | TATTCAGACTACATTAAAGGGAGTAACCTCGACGCACCTGAGCCCTATCGGATTGGACGG | 3000 |
| Dnmt1-ΔpCat | TATTCAGACTACATTAAAGGGAGTAACCTCGACGCACCTGAGCCCTATCGGATTGGACGG | 3000 |
| Dnmt1-ΔNLS | TATTCAGACTACATTAAAGGGAGTAACCTCGACGCACCTGAGCCCTATCGGATTGGACGG | 2916 |
| Dnmt1-WT | ATAAAAGAGATTCACTGTGGAAAAAAAAAGGGCAAGGTCAACGAAGCTGATATAAAACTT | 3060 |
| Dnmt1-ΔCat | ATAAAAGAGATTCACTGTGGAAAAAAAAAGGGCAAGGTCAACGAAGCTGATATAAAACTT | 3060 |
| Dnmt1-ΔpCat | ATAAAAGAGATTCACTGTGGAAAAAAAAAGGGCAAGGTCAACGAAGCTGATATAAAACTT | 3060 |
| Dnmt1-ΔNLS | ATAAAAGAGATTCACTGTGGAAAAAAAAAGGGCAAGGTCAACGAAGCTGATATAAAACTT | 2976 |
| Dnmt1-WT | AGGCTCTACAAATTTTACCGCCCTGAAAACACCCACCGATCATATAATGGTTCATACCAT | 3120 |
| Dnmt1-ΔCat | AGGCTCTACAAATTTTACCGCCCTGAAAACACCCACCGATCATATAATGGTTCATACCAT | 3120 |
| Dnmt1-ΔpCat | AGGCTCTACAAATTTTACCGCCCTGAAAACACCCACCGATCATATAATGGTTCATACCAT | 3120 |
| Dnmt1-ΔNLS | AGGCTCTACAAATTTTACCGCCCTGAAAACACCCACCGATCATATAATGGTTCATACCAT | 3036 |
| Dnmt1-WT | ACAGATATAAATATGCTTTACTGGTCCGATGAAGAAGCAGTTGTGAACTTTTCCGATGTA | 3180 |
| Dnmt1-ΔCat | ACAGATATAAATATGCTTTACTGGTCCGATGAAGAAGCAGTTGTGAACTTTTCCGATGTA | 3180 |
| Dnmt1-ΔpCat | ACAGATATAAATATGCTTTACTGGTCCGATGAAGAAGCAGTTGTGAACTTTTCCGATGTA | 3180 |
| Dnmt1-ΔNLS | ACAGATATAAATATGCTTTACTGGTCCGATGAAGAAGCAGTTGTGAACTTTTCCGATGTA | 3096 |
| Dnmt1-WT | CAGGGCAGATGTACCGTGGAATATGGAGAAGATCTGTTGGAATCTATTCAGGACTACTCA | 3240 |
| Dnmt1-ΔCat | CAGGGCAGATGTACCGTGGAATATGGAGAAGATCTGTTGGAATCTATTCAGGACTACTCA | 3240 |
| Dnmt1-ΔpCat | CAGGGCAGATGTACCGTGGAATATGGAGAAGATCTGTTGGAATCTATTCAGGACTACTCA | 3240 |
| Dnmt1-ΔNLS | CAGGGCAGATGTACCGTGGAATATGGAGAAGATCTGTTGGAATCTATTCAGGACTACTCA | 3156 |
| Dnmt1-WT | CAGGGTGGTCCCGATCGGTTCTACTTTCTCGAGGCTTACAACTCAAAGACTAAGAATTTC | 3300 |
| Dnmt1-ΔCat | CAGGGTGGTCCCGATCGGTTCTACTTTCTCGAGGCTTACAACTCAAAGACTAAGAATTTC | 3300 |

Dnmt1-ΔpCat CAGGGTGGTCCCGATCGGTTCTACTTTCTCGAGGCTTACAACTCAAAGACTAAGAATTTC 3300
Dnmt1-ΔNLS CAGGGTGGTCCCGATCGGTTCTACTTTCTCGAGGCTTACAACTCAAAGACTAAGAATTTC 3216

Dnmt1-WT GAGGATCCCCCCAACCACGCCCGAAGCCCAGGCAACAAAGGTAAAGGCAAGGGAAAGGGG 3360
Dnmt1-ΔCat GAGGATCCCCCCAACCACGCCCGAAGCCCAGGCAACAAAGGTAAAGGCAAGGGAAAGGGG 3360
Dnmt1-ΔpCat GAGGATCCCCCCAACCACGCCCGAAGCCCAGGCAACAAAGGTAAAGGCAAGGGAAAGGGG 3360
Dnmt1-ΔNLS GAGGATCCCCCCAACCACGCCCGAAGCCCAGGCAACAAAGGTAAAGGCAAGGGAAAGGGG 3276

Dnmt1-WT AAAGGCAAAGGCAAGCACCAAGTCTCAGAGCCCAAGGAGCCTGAAGCTGCTATTAAGCTG 3420
Dnmt1-ΔCat AAAGGCAAAGGCAAGCACCAAGTCTCAGAGCCCAAGGAGCCTGAAGCTGCTATTAAGCTG 3420
Dnmt1-ΔpCat AAAGGCAAAGGCAAGCACCAAGTCTCAGAGCCCAAGGAGCCTGAAGCTGCTATTAAGCTG 3420
Dnmt1-ΔNLS AAAGGCAAAGGCAAGCACCAAGTCTCAGAGCCCAAGGAGCCTGAAGCTGCTATTAAGCTG 3336

Dnmt1-WT CCCAAGCTCCGAACACTGGATGTCTTTAGTGGCTGCGGTGGATTGAGCGAAGGATTTCAT 3480
Dnmt1-ΔCat CCCAAGTAG--------------------------------------------------- 3429
Dnmt1-ΔpCat CCCAAGCTCCGAACACTGGATGTCTTTAGTGGCTGCGGTGGATTGAGCGAAGGATTTCAT 3480
Dnmt1-ΔNLS CCCAAGCTCCGAACACTGGATGTCTTTAGTGGCTGCGGTGGATTGAGCGAAGGATTTCAT 3396

Dnmt1-WT CAAGCAGGCATATCCGAAACCCTTTGGGCCATTGAAATGTGGGACCCCGCAGCACAAGCT 3540
Dnmt1-ΔCat ------------------------------------------------------------ 3429
Dnmt1-ΔpCat CAAGCAGGCATATCCGAAACCCTTTGGGCCATTGAAATGTGGGACCCCGCAGCACAAGCT 3540
Dnmt1-ΔNLS CAAGCAGGCATATCCGAAACCCTTTGGGCCATTGAAATGTGGGACCCCGCAGCACAAGCT 3456

Dnmt1-WT TTTCGCTTGAACAATCCTGGAACTACAGTCTTTACCGAAGATTGTAACGTGCTCTTGAAA 3600
Dnmt1-ΔCat ------------------------------------------------------------ 3429
Dnmt1-ΔpCat TTTCGCTTGAACAATCCTGGAACTACAGTCTTTACCGAAGATTGTAACGTGCTCTTGAAA 3600
Dnmt1-ΔNLS TTTCGCTTGAACAATCCTGGAACTACAGTCTTTACCGAAGATTGTAACGTGCTCTTGAAA 3516

Dnmt1-WT CTTGTAATGGCTGGCGAAGTCACTAACTCCCTGGGTCAACGGCTGCCACAGAAGGGTGAC 3660
Dnmt1-ΔCat ------------------------------------------------------------ 3429
Dnmt1-ΔpCat CTTGTAATGGCTGGCGAAGTCACTAACTCCCTGGGTCAACGGCTGCCACAGAAGGGTGAC 3660
Dnmt1-ΔNLS CTTGTAATGGCTGGCGAAGTCACTAACTCCCTGGGTCAACGGCTGCCACAGAAGGGTGAC 3576

Dnmt1-WT GTCGAGATGTTGTGCGGAGGTCCCCCATGTCAGGGGTTCAGCGGGATGAATAGGTTTAAC 3720
Dnmt1-ΔCat ------------------------------------------------------------ 3429
Dnmt1-ΔpCat GTCGAGATGTTGTGCGGAGGTCCCGGCAGCCAGGGGTTCAGCGGGATGAATAGGTTTAAC 3720
Dnmt1-ΔNLS GTCGAGATGTTGTGCGGAGGTCCCCCATGTCAGGGGTTCAGCGGGATGAATAGGTTTAAC 3636

Dnmt1-WT TCACGGACATACTCCAAGTTCAAGAACAGTCTCGTGGTCAGTTTCTTGTCTTACTGCGAT 3780
Dnmt1-ΔCat ------------------------------------------------------------ 3429
Dnmt1-ΔpCat TCACGGACATACTCCAAGTTCAAGAACAGTCTCGTGGTCAGTTTCTTGTCTTACTGCGAT 3780
Dnmt1-ΔNLS TCACGGACATACTCCAAGTTCAAGAACAGTCTCGTGGTCAGTTTCTTGTCTTACTGCGAT 3696

Dnmt1-WT TATTATAGACCCCGATTTTTTTTGCTTGAGAACGTAAGAAATTTCGTAAGTTACCGGAGG 3840
Dnmt1-ΔCat ------------------------------------------------------------ 3429
Dnmt1-ΔpCat TATTATAGACCCCGATTTTTTTTGCTTGAGAACGTAAGAAATTTCGTAAGTTACCGGAGG 3840
Dnmt1-ΔNLS TATTATAGACCCCGATTTTTTTTGCTTGAGAACGTAAGAAATTTCGTAAGTTACCGGAGG 3756

Dnmt1-WT AGCATGGTGCTGAAGCTGACCTTGAGATGCCTCGTGAGGATGGGGTACCAATGCACCTTT 3900
Dnmt1-ΔCat ------------------------------------------------------------ 3429
Dnmt1-ΔpCat AGCATGGTGCTGAAGCTGACCTTGAGATGCCTCGTGAGGATGGGGTACCAATGCACCTTT 3900
Dnmt1-ΔNLS AGCATGGTGCTGAAGCTGACCTTGAGATGCCTCGTGAGGATGGGGTACCAATGCACCTTT 3816

Dnmt1-WT GGGGTACTGCAGGCCGGACAGTATGGAGTGGCACAGACTCGCCGCCGAGCAATAATCCTT 3960
Dnmt1-ΔCat ------------------------------------------------------------ 3429
Dnmt1-ΔpCat GGGGTACTGCAGGCCGGACAGTATGGAGTGGCACAGACTCGCCGCCGAGCAATAATCCTT 3960
Dnmt1-ΔNLS GGGGTACTGCAGGCCGGACAGTATGGAGTGGCACAGACTCGCCGCCGAGCAATAATCCTT 3876

Dnmt1-WT GCTGCAGCTCCAGGTGAGAAACTTCCTCTGTTCCCCGAACCACTTCACGTTTTTGCACCA 4020
Dnmt1-ΔCat ------------------------------------------------------------ 3429
Dnmt1-ΔpCat GCTGCAGCTCCAGGTGAGAAACTTCCTCTGTTCCCCGAACCACTTCACGTTTTTGCACCA 4020
Dnmt1-ΔNLS GCTGCAGCTCCAGGTGAGAAACTTCCTCTGTTCCCCGAACCACTTCACGTTTTTGCACCA 3936

Dnmt1-WT AGAGCTTGCCAACTGTCAGTCGTTGTTGATGACAAAAAGTTCGTCTCTAACATCACAAGA 4080
Dnmt1-ΔCat ------------------------------------------------------------ 3429
Dnmt1-ΔpCat AGAGCTTGCCAACTGTCAGTCGTTGTTGATGACAAAAAGTTCGTCTCTAACATCACAAGA 4080
Dnmt1-ΔNLS AGAGCTTGCCAACTGTCAGTCGTTGTTGATGACAAAAAGTTCGTCTCTAACATCACAAGA 3996

Dnmt1-WT CTTTCCTCTGGACCATTCCGCACCATAACAGTCCGCGACACCATGTCAGATTTGCCAGAA 4140
Dnmt1-ΔCat ------------------------------------------------------------ 3429
Dnmt1-ΔpCat CTTTCCTCTGGACCATTCCGCACCATAACAGTCCGCGACACCATGTCAGATTTGCCAGAA 4140
Dnmt1-ΔNLS CTTTCCTCTGGACCATTCCGCACCATAACAGTCCGCGACACCATGTCAGATTTGCCAGAA 4056

Dnmt1-WT ATACAGAACGGGGCATCAAACTCCGAGATCCCATACAATGGGGAGCCACTTTCATGGTTT 4200
Dnmt1-ΔCat ------------------------------------------------------------ 3429
Dnmt1-ΔpCat ATACAGAACGGGGCATCAAACTCCGAGATCCCATACAATGGGGAGCCACTTTCATGGTTT 4200
Dnmt1-ΔNLS ATACAGAACGGGGCATCAAACTCCGAGATCCCATACAATGGGGAGCCACTTTCATGGTTT 4116

```
Dnmt1-WT      CAGCGACAACTGAGGGGGTCCCACTATCAGCCAATACTCCGCGACCATATCTGTAAAGAT   4260
Dnmt1-ΔCat    ------------------------------------------------------------   3429
Dnmt1-ΔpCat   CAGCGACAACTGAGGGGGTCCCACTATCAGCCAATACTCCGCGACCATATCTGTAAAGAT   4260
Dnmt1-ΔNLS    CAGCGACAACTGAGGGGGTCCCACTATCAGCCAATACTCCGCGACCATATCTGTAAAGAT   4176

Dnmt1-WT      ATGTCTCCATTGGTAGCAGCCCGGATGCGACACATACCCCTTTTTCCAGGATCAGACTGG   4320
Dnmt1-ΔCat    ------------------------------------------------------------   3429
Dnmt1-ΔpCat   ATGTCTCCATTGGTAGCAGCCCGGATGCGACACATACCCCTTTTTCCAGGATCAGACTGG   4320
Dnmt1-ΔNLS    ATGTCTCCATTGGTAGCAGCCCGGATGCGACACATACCCCTTTTTCCAGGATCAGACTGG   4236

Dnmt1-WT      CGGGACTTGCCTAACATACAGGTAAGGCTCGGGGACGGAGTGATAGCACACAAACTCCAA   4380
Dnmt1-ΔCat    ------------------------------------------------------------   3429
Dnmt1-ΔpCat   CGGGACTTGCCTAACATACAGGTAAGGCTCGGGGACGGAGTGATAGCACACAAACTCCAA   4380
Dnmt1-ΔNLS    CGGGACTTGCCTAACATACAGGTAAGGCTCGGGGACGGAGTGATAGCACACAAACTCCAA   4296

Dnmt1-WT      TATACTTTCCATGACGTAAAGAATGGCTACTCTTCCACAGGAGCTTTGCGAGGTGTCTGC   4440
Dnmt1-ΔCat    ------------------------------------------------------------   3429
Dnmt1-ΔpCat   TATACTTTCCATGACGTAAAGAATGGCTACTCTTCCACAGGAGCTTTGCGAGGTGTCTGC   4440
Dnmt1-ΔNLS    TATACTTTCCATGACGTAAAGAATGGCTACTCTTCCACAGGAGCTTTGCGAGGTGTCTGC   4356

Dnmt1-WT      AGTTGCGCCGAAGGAAAGGCTTGCGATCCAGAATCCCGCCAATTTAGTACCCTTATACCC   4500
Dnmt1-ΔCat    ------------------------------------------------------------   3429
Dnmt1-ΔpCat   AGTTGCGCCGAAGGAAAGGCTTGCGATCCAGAATCCCGCCAATTTAGTACCCTTATACCC   4500
Dnmt1-ΔNLS    AGTTGCGCCGAAGGAAAGGCTTGCGATCCAGAATCCCGCCAATTTAGTACCCTTATACCC   4416

Dnmt1-WT      TGGTGCCTTCCACATACAGGAAATCGACACAACCACTGGGCCGGTCTCTATGGAAGGCTC   4560
Dnmt1-ΔCat    ------------------------------------------------------------   3429
Dnmt1-ΔpCat   TGGTGCCTTCCACATACAGGAAATCGACACAACCACTGGGCCGGTCTCTATGGAAGGCTC   4560
Dnmt1-ΔNLS    TGGTGCCTTCCACATACAGGAAATCGACACAACCACTGGGCCGGTCTCTATGGAAGGCTC   4476

Dnmt1-WT      GAGTGGGATGGATTTTTCTCTACTACTGTTACAAATCCTGAACCCATGGGCAAGCAAGGT   4620
Dnmt1-ΔCat    ------------------------------------------------------------   3429
Dnmt1-ΔpCat   GAGTGGGATGGATTTTTCTCTACTACTGTTACAAATCCTGAACCCATGGGCAAGCAAGGT   4620
Dnmt1-ΔNLS    GAGTGGGATGGATTTTTCTCTACTACTGTTACAAATCCTGAACCCATGGGCAAGCAAGGT   4536

Dnmt1-WT      CGCGTTCTCCATCCAGAACAACACCGAGTGGTGTCAGTCCGGGAATGCGCTCGGTCCCAA   4680
Dnmt1-ΔCat    ------------------------------------------------------------   3429
Dnmt1-ΔpCat   CGCGTTCTCCATCCAGAACAACACCGAGTGGTGTCAGTCCGGGAATGCGCTCGGTCCCAA   4680
Dnmt1-ΔNLS    CGCGTTCTCCATCCAGAACAACACCGAGTGGTGTCAGTCCGGGAATGCGCTCGGTCCCAA   4596

Dnmt1-WT      GGTTTCCCTGACTCATACAGATTTTTTGGAAACATTCTCGACCGCCATCGCCAGGTTGGT   4740
Dnmt1-ΔCat    ------------------------------------------------------------   3429
Dnmt1-ΔpCat   GGTTTCCCTGACTCATACAGATTTTTTGGAAACATTCTCGACCGCCATCGCCAGGTTGGT   4740
Dnmt1-ΔNLS    GGTTTCCCTGACTCATACAGATTTTTTGGAAACATTCTCGACCGCCATCGCCAGGTTGGT   4656

Dnmt1-WT      AACGCCGTGCCACCCCCCCTCGCAAAAGCTATAGGGCTTGAGATCAAGTTGTGCTTGTTG   4800
Dnmt1-ΔCat    ------------------------------------------------------------   3429
Dnmt1-ΔpCat   AACGCCGTGCCACCCCCCCTCGCAAAAGCTATAGGGCTTGAGATCAAGTTGTGCTTGTTG   4800
Dnmt1-ΔNLS    AACGCCGTGCCACCCCCCCTCGCAAAAGCTATAGGGCTTGAGATCAAGTTGTGCTTGTTG   4716

Dnmt1-WT      AGTTCAGCAAGGGAATCAGCTTCAGCAGCCGTTAAGGCAAAGGAGGAAGCTGCCACTAAG   4860
Dnmt1-ΔCat    ------------------------------------------------------------   3429
Dnmt1-ΔpCat   AGTTCAGCAAGGGAATCAGCTTCAGCAGCCGTTAAGGCAAAGGAGGAAGCTGCCACTAAG   4860
Dnmt1-ΔNLS    AGTTCAGCAAGGGAATCAGCTTCAGCAGCCGTTAAGGCAAAGGAGGAAGCTGCCACTAAG   4776

Dnmt1-WT      GATTAG      4866
Dnmt1-ΔCat    ------      3429
Dnmt1-ΔpCat   GATTAG      4866
Dnmt1-ΔNLS    GATTAG      4782
```

#### 4 Supplementary Data Legend

**Supplementary data 1:**

List of proteins being co-immunoprecipitated with DNMT1 using an antibody targeting DNMT1 in murine brain tissue lysates (3.5 months), which show a fold change of at least two, identified by mass spectrometry. N = 4 independent experiments. Two-tailed Student's t-test, ** $p < 0.01$, *** $p < 0.001$, **** $p < 0.0001$.

**Supplementary data 2:**

List of proteins being co-immunoprecipitated with DNMT1 using an antibody targeting DNMT1 in N2a cell lysates, which show a fold change of at least two, identified by mass spectrometry. N = 4 independent experiments. Two-tailed Student's t-test, ** $p < 0.01$, *** $p < 0.001$, **** $p < 0.0001$.

#### 5 Supplementary Methods

##### Microscopy (Fixed cells)

For DNMT1 subcellular localization and colocalization with DOCK7, imaging was performed using a STED microscope (TCS SP8, Leica, Germany) with a 93x/NA 1.3, HC PL APO CS2 glycerol immersion objective. Excitation was achieved using a pulsed white light laser (WLL) to stimulate the respective fluorophores at 405 nm (DAPI, nuclei, 470 nm emission), 554 nm (Cy3, DOCK7, 566 nm emission), and 633 nm (Cy5, DNMT1, 670 nm emission). The emissions were detected via HyD time-gated mode (0.5-6.0 ns), and STED depletion was applied at 775 nm. Images were acquired at 600 Hz with a voxel size of 0.05 × 0.05 × 0.18 $\mu$m and 512 × 512 px resolution. Deconvolution was executed using *Imaris 10* software (Oxford Instruments, UK). For DNMT1/DOCK7 colocalization with mitochondria, the excitation was set at 405 nm (DAPI, nuclei, 470 nm emission), 499 nm (Alexa488, DNMT1/DOCK7, 525 nm emission), and 649 nm (Cy5, MitoTracker™ Deep Red FM, 670 nm emission). The other parameters were applied equally as described above. The imaging was executed using the *LAS X Core* software and *LAS X Navigator* (Leica, Germany) by the Core Facility for 3D Superresolution (RWTH Aachen University, Germany).

The imaging of CRISPRi-Ctrl/Dnmt1-GFP-positive neurons and immunocytochemically stained brain sections was executed using the FV1000 and FV4000 confocal laser scanning microscopes (Olympus, Japan).

Imaging of immunocytochemically stained acetylated tubulin (AcTUB) in fixed cells was performed using the BZ-X810 fluorescence microscope (Keyence, Japan) with a Plan Apochromat 40x/NA 0.95 M25 objective. The fluorescence channels were adjusted as follows: for the Cy5-conjugated AcTUB (excitation 640 nm, emission 670 nm), for Alexa555-labeled control siRNA (TRITC, excitation 555 nm, emission 578 nm), for Alexa488-conjugated βIII-tubulin (FITC, excitation 488 nm, emission 520 nm), and for DAPI (excitation 405 nm, emission 470 nm). All imaging procedures were conducted with *High Resolution* settings and *Low Photobleach* mode, without z-stacks or post-processing using the *VK-X3000 Viewer* software (Keyence, Japan).

Fixed cells following immunocytochemical staining were imaged using the DMi8 inverted microscope combined with the THUNDER® imaging platform (Leica, Germany). Relative Focus Correction (RFC) was applied for all multichannel imaging workflows. All images were post-processed by Instant Computational Clearing (ICC) without binning and without z-stacks. To validate the CRISPRi-Dnmt1-GFP construct, the HC PL FLUOTAR L 40x/0.60 DRY objective was used (pixel size: 0.16 × 0.16 $\mu$m; xy resolution: 0.559 × 0.559 $\mu$m). For morphological analysis, either the HC PL FLUOTAR L 20x/0.40 DRY objective (used for cortical neurons at E14.5 + 2 DIV; pixel size: 0.32 × 0.32 $\mu$m; xy resolution: 665.28 × 665.28 $\mu$m) or the HC PL FLUOTAR L 40x/0.60 DRY objective (used for N2a cells) was employed. The HC PL FLUOTAR L 20x/0.40 DRY objective was also used for RG108 validation and SMI-312 immunocytochemical staining. For localization studies of mNG-DNMT1 in N2a cells, the HC PL APO 40x/1.30 OIL objective was used (pixel size: 0.16 × 0.16 $\mu$m; xy resolution: 0.258 × 0.258 $\mu$m). DAPI filter settings (excitation: 395 nm; emission: 434 nm) were used to visualize nuclei, and the brightfield channel provided a general overview of the cells. TRITC filter settings (excitation: 550 nm; emission: 595 nm) were applied for imaging Alexa555-labeled control siRNA or CAG-mSc expression. FITC filter settings (excitation: 470 nm; emission: 512 nm) were used for detection of Alexa488-labeled control siRNA, various CAG-mNG-*Dnmt1* constructs, or CRISPRi-Ctrl/Dnmt1-GFP expression. Far-red filter settings (excitation: 640 nm; emission: 705 nm) were employed to visualize Cy5-labeled control siRNA, DNMT1, 5mC, or βIII-tubulin. All images were acquired at a resolution of 2048 × 2048 pixels, with a scanning speed of 216 MHz applied throughout all imaging procedures. Imaging was executed using the *LAS X Core* software (Leica, Germany).

**Microscopy (Live cell imaging)**

Live cell imaging of mitochondrial accumulation was performed using a STED microscope (TCS SP8, Leica, Germany), including an incubation unit ("BOX", "ICE CUBE", "BRICK"; Life Imaging Services, Switzerland), maintaining a constant atmosphere of 37 °C and 5% $CO_2$. A 63x/NA 1.4, HC PL APO CS2 oil immersion objective was used. Excitation was set at 499 nm for Alexa488-conjugated control siRNA (525 nm emission) and at 649 nm for MitoTracker™ Deep Red FM-positive organelles (670 nm emission) using a pulsed white light laser (WLL). N2a cells were localized via the *LAS X Navigator* (Leica, Germany), and sequential z-stacks were recorded every 2 min for 5 h. Images were acquired at 600 Hz (bidirectional) with a voxel size of 0.36 × 0.36 × 0.3 $\mu$m (x, y, z) and 512 × 512 px resolution. Imaging was performed with the *LAS X Core* software (Leica, Germany) at the Core Facility for 3D Superresolution (RWTH Aachen University, Germany).

Live cell imaging of mitochondrial trafficking and EB3 comet dynamics were conducted using a spinning disk confocal microscope (Dragonfly, Oxford Instruments Andor, UK) with a 100x/NA 1.49, CFI SR HP Apo TIRF oil immersion objective and an integrated incubation unit, maintaining a constant atmosphere of 37 °C and 5% $CO_2$. P0 + 4 DIV cortical neurons were checked beforehand for Alexa555-positive control siRNA using TRITC filter settings (excitation 521 nm, emission 568 nm). MT-GFP-positive mitochondria were imaged using FITC filter settings (excitation 488 nm, emission 525 nm) without z-stacks. In experiments involving mNG-*Dnmt1* plasmids, MT-DsRed plasmid, and Cy5-labeled control siRNA, mitochondria were visualized using TRITC filter settings (excitation 594 nm, emission 578 nm), mNG-DNMT1 expression via FITC filter settings (excitation 488 nm, emission 525 nm), and control siRNA via Cy5 filter settings (excitation 637 nm, emission 670 nm). One frame was recorded every 3 sec for 5 min (100 frames in total) during time-lapse imaging, with a pixel size of 6.5 × 6.5 $\mu$m (x, y) and 512 × 512 px resolution. Scanning speed was set at 100 MHz. The imaging and further deconvolution were applied using the *Fusion* software (Oxford Instruments Andor, UK).

The TMRE and FCCP validation experiments were imaged using the BZ-X810 (Keyence, Japan) and DMi8 with a THUNDER® imager unit (Leica, Germany) fluorescence microscope with the respective integrated incubation units, which maintain a constant atmosphere of 37 °C and 5% $CO_2$. TMRE fluorescence was detected using the TRITC filter settings (excitation 550 nm, emission 578 nm), and Alexa488-labeled control siRNA was visualized using the FITC filter settings (excitation 488 nm, emission 520 nm). Brightfield imaging was used for general cell visualization and overview. Each image was captured with 5 z-stacks (0.494 $\mu$m z-resolution). Images derived from the BZ-X810 (Keyence, Japan) using the 40x/NA 0.95 Plan Apochromat objective were acquired with *High Resolution* settings, *Low Photobleach* mode, and without post-processing. Images derived from the DMi8 (Leica, Germany) were generated using the 40x/0.60 dry objective (2048 × 2048 px) and instant computational clearing (ICC) was conducted as post-processing. The scanning speed was set to 216 MHz. The pixel size was 0.16 × 0.16 $\mu$m (x, y) and the resolution 0.559 x 0.559 $\mu$m. Imaging was performed using *LAS X Core* software (Leica, Germany) and the *VK-X3000 Viewer* (Keyence, Japan).

### Supplementary Figure S1:

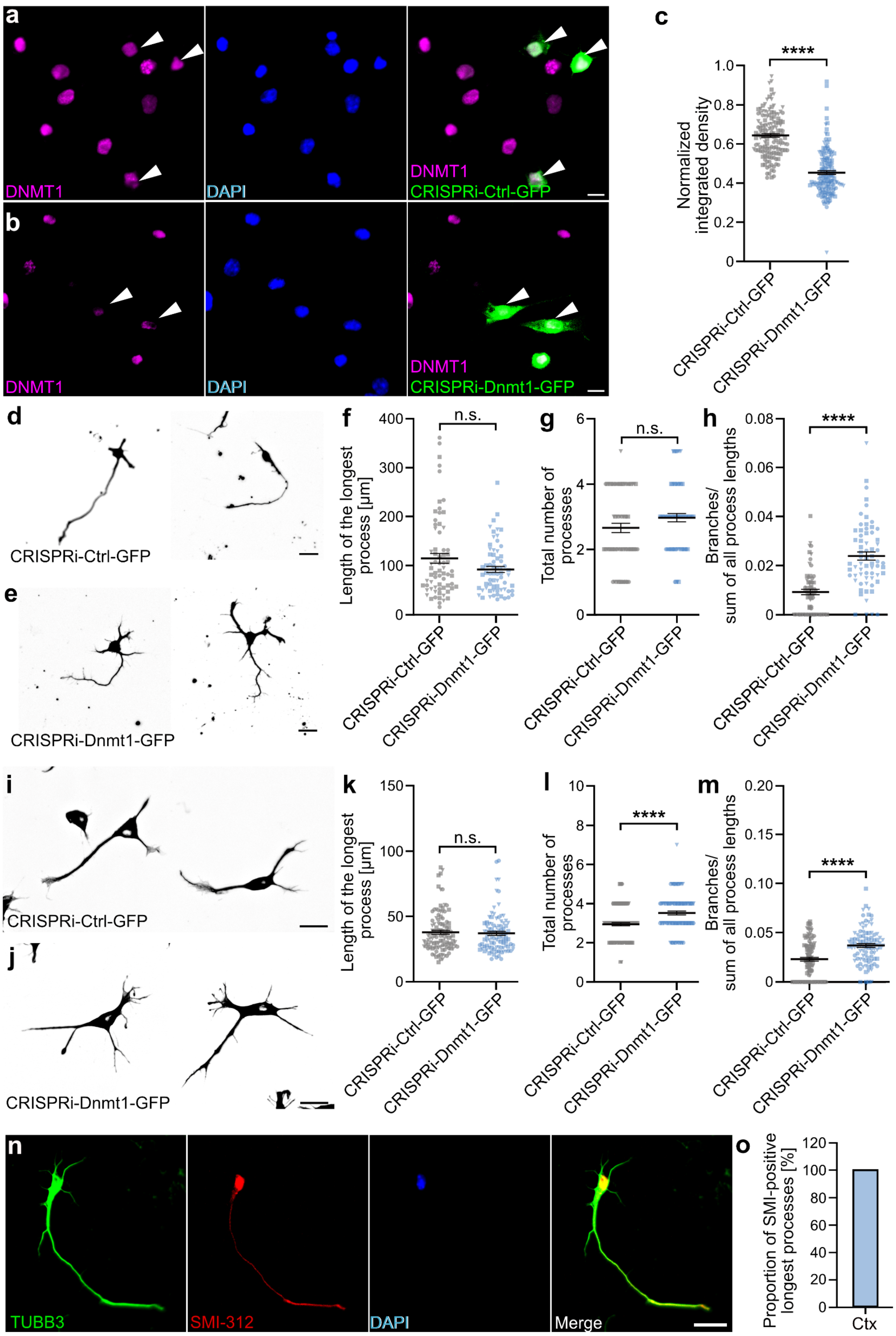

### Supplementary Figure S2:

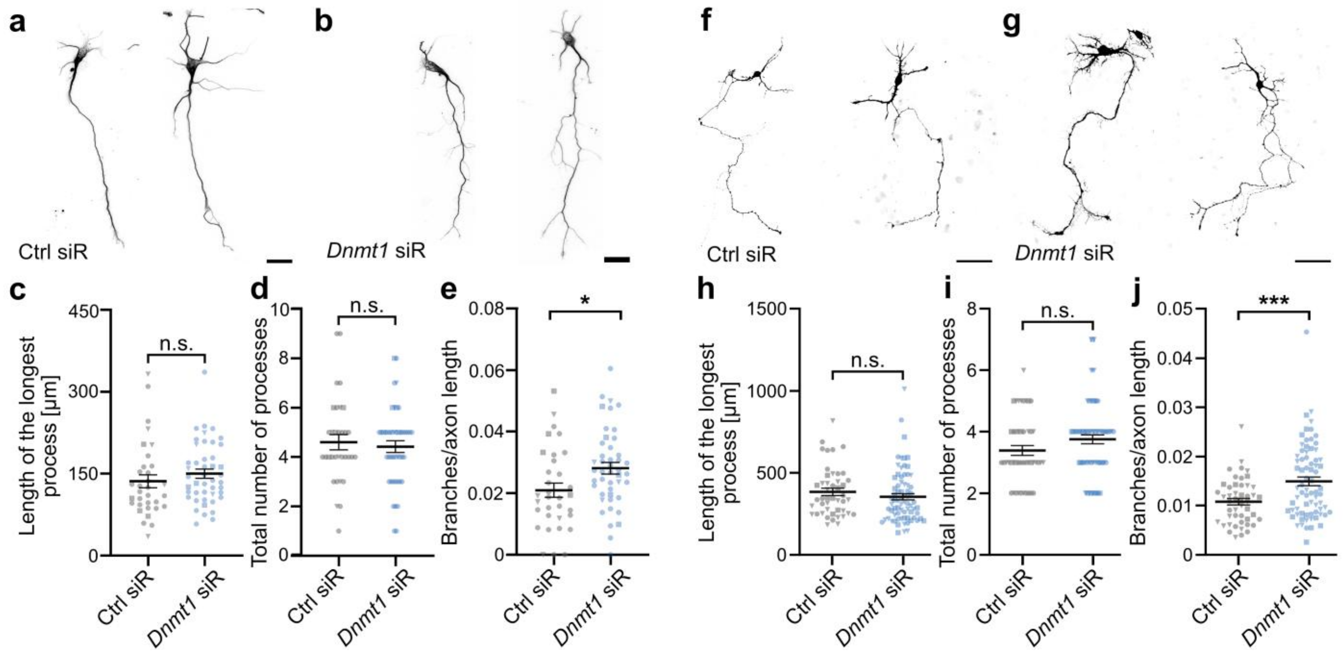

### Supplementary Figure S3:

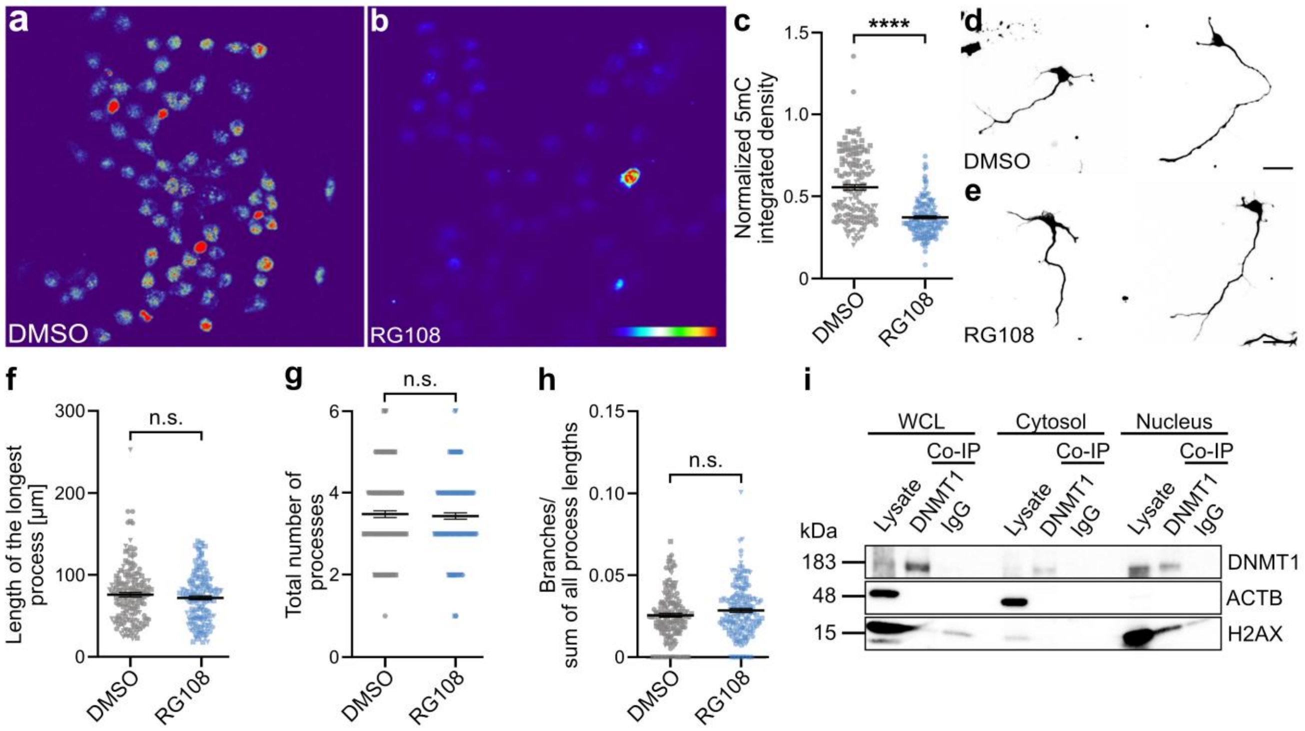

### Supplementary Figure S4:

**a**

| ID | Name | Abb. |
|---|---|---|
| Q8R1A4 | Dedicator of cytokinesis protein 7 | Dock7 |
| Q9CYZ2 | Tumor protein D54 | Tpd52l2 |
| P27546 | Microtubule-associated protein 4 | Map4 |
| P33173 | Kinesin-like protein KIF1A | Kif1a |
| A2AG50 | MAP7 domain-containing protein 2 | Map7d2 |
| P20357 | Microtubule-associated protein 2 | Map2 |
| Q61768 | Kinesin-1 heavy chain | Kif5b |
| Q8VDR9 | Dedicator of cytokinesis protein 6 | Dock6 |
| Q9QWT9 | Kinesin-like protein KIFC1 | Kifc1 |

**b**

| ID | Name | Abb. |
|---|---|---|
| Q8BX10 | Mitochondrial serine/threonine protein phosphatase | Pgam5 |
| P58281 | Mitochondrial dynamin like GTPase | Opa1 |
| P35486 | Pyruvate dehydrogenase E1 subunit alpha 1 | Pdha1 |
| P26443 | Glutamate dehydrogenase 1 | Glud1 |
| P47738 | Aldehyde dehydrogenase, mitochondrial | Aldh2 |
| Q3U2A8 | Valyl-tRNA synthetase 2 | Vars2 |
| Q3UFY8 | tRNA methyltransferase 10C | Trmt10c |
| P28352 | Apurinic/Apyrimidinic endodeoxyribonuclease 1 | Apex1 |
| P61922 | 4-Aminobutyrate aminotransferase | Abat |
| Q9D2G2 | Dihydrolipoamide S-succinyltransferase | Dlst |

**c**

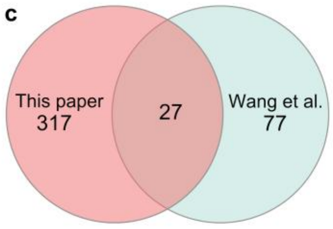

**d** **e**

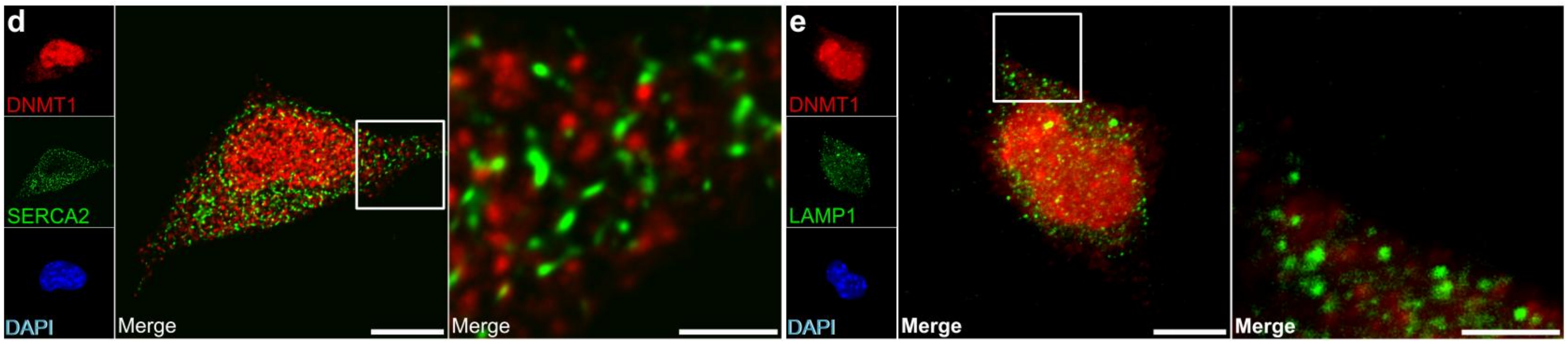

### Supplementary Figure S5:

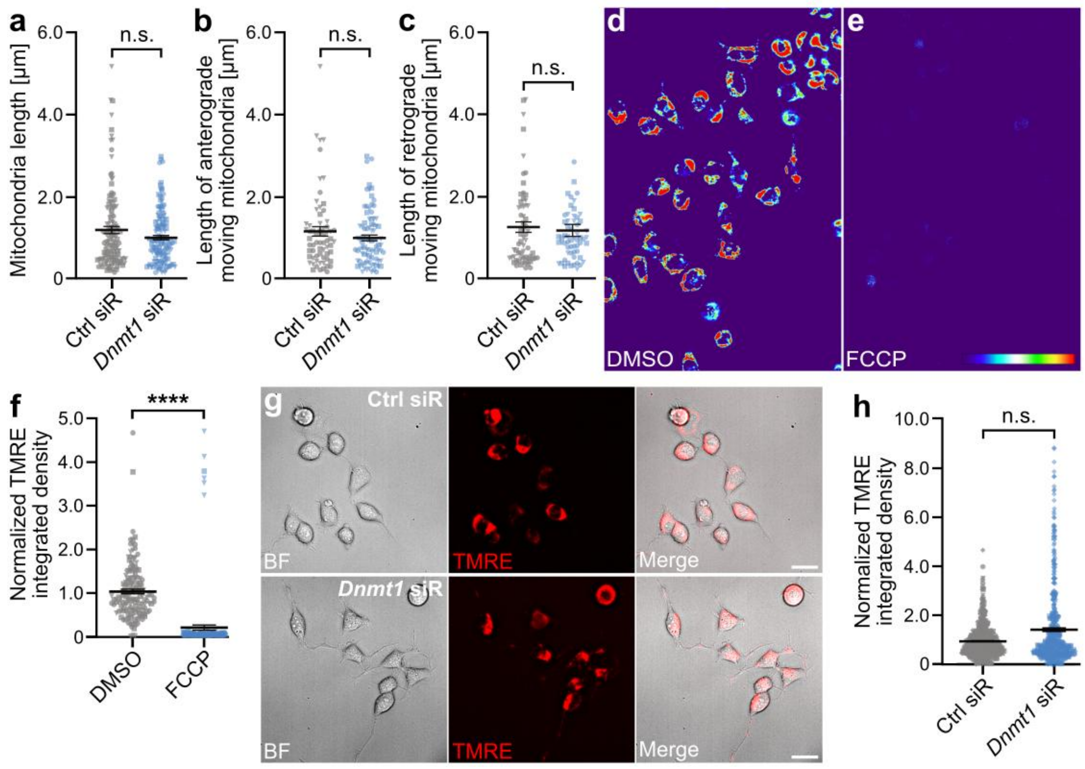

### Supplementary Figure S6:

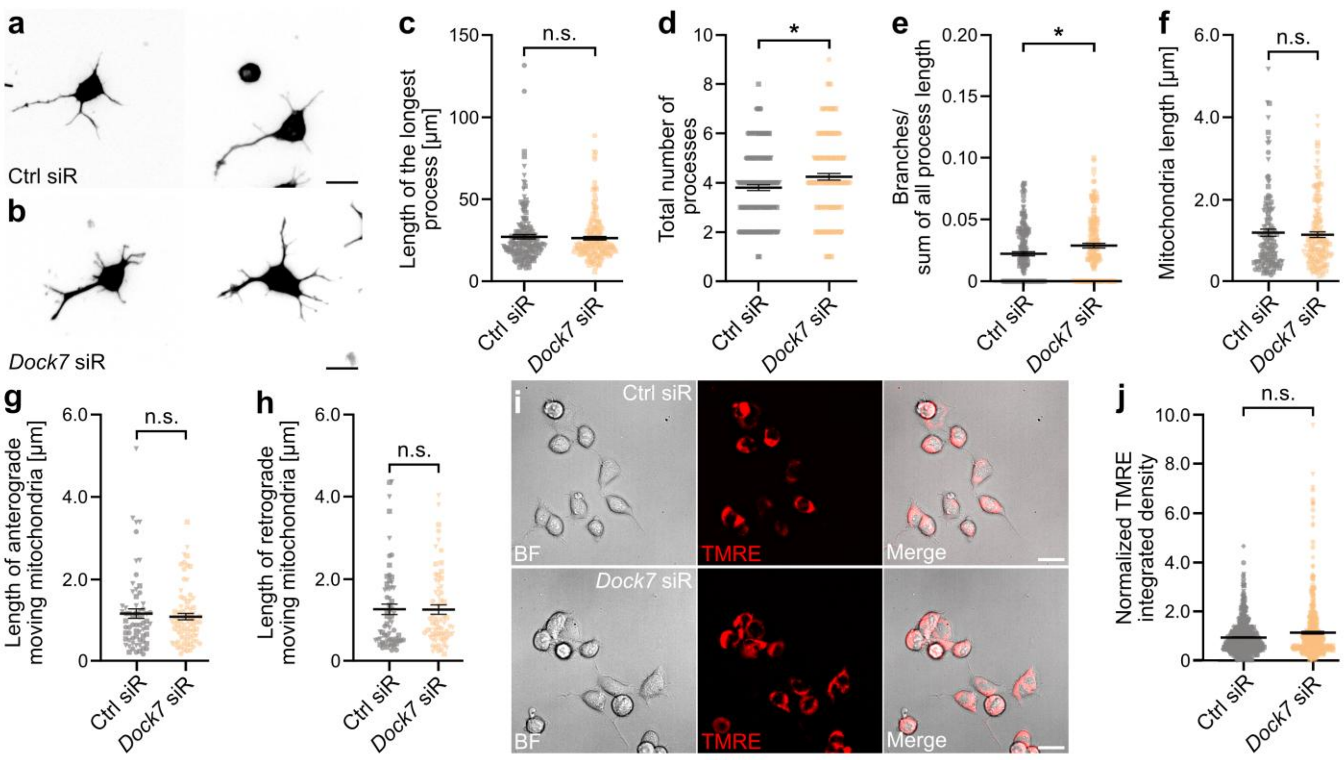

### Supplementary Figure S7:

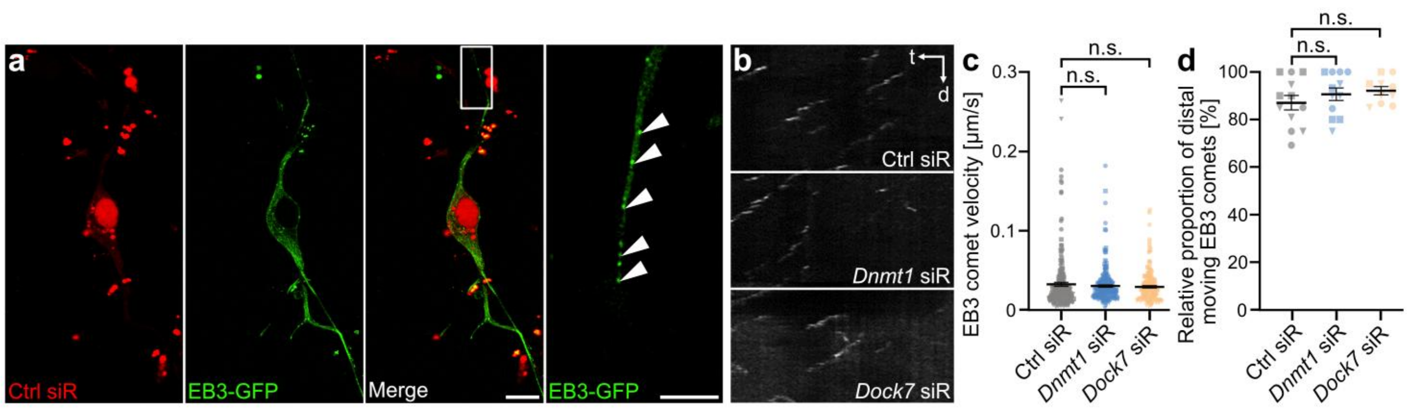
